# Decoding polyubiquitin regulation of K_V_7. 1 functional expression with engineered linkage-selective deubiquitinases

**DOI:** 10.1101/2024.09.17.613539

**Authors:** Sri Karthika Shanmugam, Scott A. Kanner, Xinle Zou, Enoch Amarh, Papiya Choudhury, Rajesh Soni, Robert S. Kass, Henry M. Colecraft

**Author notes:** Correspondence: Henry M. Colecraft Department of Physiology and Cellular Biophysics Columbia University, College of Physicians and Surgeons 504 Russ Berrie Pavilion 1150 St. Nicholas Avenue New York, NY 10032. these authors contributed equally to the work (Sri Karthika Shanmugam, Scott A. Kanner).

## Abstract

Protein posttranslational modification with distinct polyubiquitin linkage chains is a critical component of the ‘ubiquitin code’ that universally regulates protein expression and function to control biology. Functional consequences of diverse polyubiquitin linkages on proteins are mostly unknown, with progress hindered by a lack of methods to specifically tune polyubiquitin linkages on individual proteins in live cells. Here, we bridge this gap by exploiting deubiquitinases (DUBs) with preferences for hydrolyzing different polyubiquitin linkages: OTUD1 - K63; OTUD4 - K48; Cezanne - K11; TRABID - K29/K33; and USP21 - non-specific. We developed a suite of engineered deubiquitinases (enDUBs) comprised of DUB catalytic domains fused to a GFP-targeted nanobody and used them to investigate polyubiquitin linkage regulation of an ion channel, YFP-KCNQ1. Mass spectrometry of YFP-KCNQ1 expressed in HEK293 cells indicated channel polyubiquitination with K48 (72%) and K63 (24%) linkages being dominant. NEDD4-2 and ITCH both decreased KCNQ1 functional expression but with distinctive polyubiquitination signatures. All enDUBs reduced KCNQ1 ubiquitination but yielded unique effects on channel expression, surface density, ionic currents, and subcellular localization. The pattern of outcomes indicates K11, K29/K33, and K63 chains mediate net KCNQ1-YFP intracellular retention, but achieved in different ways: K11 promotes ER retention/degradation, enhances endocytosis, and reduces recycling; K29/K33 promotes ER retention/degradation; K63 enhances endocytosis and reduces recycling. The pattern of enDUB effects on KCNQ1-YFP differed in cardiomyocytes, emphasizing ubiquitin code mutability. Surprisingly, enDUB-O4 decreased KCNQ1-YFP surface density suggesting a role for K48 in forward trafficking. Lastly, linkage-selective enDUBs displayed varying capabilities to rescue distinct trafficking-deficient long QT syndrome type 1 mutations. The results reveal distinct polyubiquitin chains control different aspects of KCNQ1 functional expression, demonstrate ubiquitin code plasticity, and introduce linkage-selective enDUBs as a potent tool to help demystify the polyubiquitin code.

## Introduction

Ubiquitin is a pervasive regulator of protein expression and cell signaling pathways and, thereby, controls all aspects of biology. Protein ubiquitination is accomplished through the sequential activities of three enzyme classes— E1 enzymes catalyze ATP-dependent activation and transfer of ubiquitin to E2 conjugating enzymes from where it is covalently attached most frequently to the ε-amino group of substrate lysine residues through the activity of E3 ubiquitin ligases (Komander and Rape, 2012; Schulman and Harper, 2009). The human genome encodes 2 E1, ∼40 E2, and >600 E3 enzymes. Ubiquitin itself has seven internal lysine residues (K6, K11, K27, K29, K33, K48, and K63) which, along with its N-terminal M1 residue, can be ubiquitinated to produce a variety of polyubiquitin chains with distinctive structures (Akutsu et al., 2016; French et al., 2021). The capacity to produce specific polyubiquitin linkages is encoded within particular E2 and E3 ligases. Individual proteins may be modified by monoubiquitination, multi-monoubiquitination, homotypic polyubiquitination (one linkage type), and/or heterotypic polyubiquitination with mixed and/or branched chains (Akutsu et al., 2016; Komander and Rape, 2012; Kwon and Ciechanover, 2017; Yau and Rape, 2016). The distinctive topological features of diverse ubiquitin/polyubiquitin modifications are decoded by various effector proteins with domains and motifs that recognize specific ubiquitin configurations to yield discrete functional outcomes (Dikic et al., 2009; Hicke et al., 2005; Hofmann and Falquet, 2001). Protein ubiquitination is dynamically countered by ∼100 deubiquitinases (DUBs) which hydrolyze ubiquitin chains in distinctive ways (Mevissen and Komander, 2017). Altogether, the system of ubiquitin writers (E1, E2 and E3 ligases), readers (effectors), editors, and erasers (DUBs), together with the multiplicity of possible ubiquitin modification types on proteins, constitute a complex and versatile ‘ubiquitin code’ that remains largely enigmatic despite decades of research (Dikic and Schulman, 2023; Komander and Rape, 2012; Kwon and Ciechanover, 2017; Yau and Rape, 2016). The scope of the problem is colossal due to the prevalence of ubiquitination, the uniqueness of distinct cellular contexts, and the likely substrate-specific nature of functional outcomes.

Ubiquitin proteomics methods indicate a substantial fraction of cellular total ubiquitin (10-20%) is incorporated into polyubiquitin chains with K48 and K63 typically being most abundant (Swatek et al., 2019; Xu et al., 2009). A key aspect of deciphering the ubiquitin code is identifying the functional consequences of distinct polyubiquitin chains on individual proteins. In this regard, K48 polyubiquitin chains are canonically associated with protein degradation via the proteasome, the original function ascribed to protein ubiquitination (Chau et al., 1989; Clague and Urbe, 2010). K63 chains, by contrast, have been associated with diverse non-degradative functions including: activation of IκB kinase, a step necessary for triggering NFκB signaling (Deng et al., 2000; Wang et al., 2001); mediating post-replicative DNA repair via attachment to proliferating cell nuclear antigen (Hoege et al., 2002); and directing sorting of epidermal growth factor receptors (EGFR) from early to late endosomes (Huang et al., 2013). With respect to more atypical chains, examples include: the anaphase-promoting complex (APC/C) which coordinates mitosis by catalyzing K11 polyubiquitination of cell-cycle regulators that marks them for proteasomal degradation (Jin et al., 2008; Song and Rape, 2010); PARKIN-mediated K6 polyubiquitination of mitochondrial outer membrane proteins is implicated in mitophagy (Ordureau et al., 2015); and K27/K29 polyubiquitination of stimulator of interferon genes (STING) mediates STING translocation from the endoplasmic reticulum (ER) to Golgi, an obligatory step for its mechanism of action (Kong et al., 2023). These insights into functions of distinct polyubiquitin chains have been obtained through combinations of elegant biochemical studies, mass spectrometry, and evaluating the impact of K-to-R ubiquitin mutants in cells. The cell-free biochemical approaches are substrate-specific, not generalizable, and low throughput, while the use of K-to-R ubiquitin mutants yield global changes in polyubiquitination that may complicate data interpretation (Xu et al., 2009). A generalizable method that permits substrate-selective, linkage-selective hydrolysis of polyubiquitin chains from target proteins in live cells is necessary for deciphering the mysterious ubiquitin code, but is currently lacking.

A potential solution is offered by the distinctive properties of DUBs which are classified into six families – ubiquitin-specific proteases (USPs), ovarian tumor proteases (OTUs), ubiquitin C-terminal hydrolases (UCHs), Josephin family, motif interacting with ubiquitin (MIU)-containing novel DUB family (MINDYs), and the JAB1/MPN/MOV34 metalloprotease DUBs (JAMMs) (Clague et al., 2019; Komander et al., 2009). Interestingly, certain DUBs display preferential hydrolysis of particular ubiquitin linkages in polyubiquitin chains. In particular, distinct OTU family members display an impressive polyubiquitin-linkage specificity *in vitro* that is encoded within discrete catalytic domains (Mevissen et al., 2013; Mevissen and Komander, 2017). The properties of linkage-selective DUBs have been exploited for ubiquitin chain restriction analyses of ubiquitinated substrates *in vitro* (Hospenthal et al., 2015). Whether catalytic domains of linkage-selective DUBs can be exploited to hydrolyze polyubiquitin chains in a substrate-specific manner within live cells has not been systematically explored.

Cell surface ion channels are a specialized class of integral membrane proteins that are present in all organs, tissues, and cell types and are necessary for myriad biological processes (Subramanyam and Colecraft, 2015). The precise mechanisms regulating their stability and trafficking to and away from the cell surface through various intracellular compartments are largely mysterious, but known to be influenced by ubiquitination (Abriel and Staub, 2005; Rotin and Staub, 2011). Many devastating ion channelopathies arise from mutations in ion channels that adversely impact their surface density, highlighting the importance of understanding fundamental ion channel trafficking mechanisms (Curran and Mohler, 2015; Harraz and Delpire, 2024). Linkage-selective polyubiquitination is a potentially critical, but under-appreciated determinant of ion channel trafficking among distinct subcellular compartments. These general concepts are exemplified by KCNQ1 (Kv7.1/Q1) which together with auxiliary KCNE1 subunits generates the slow delayed rectifier current, *I*_Ks_, necessary for proper repolarization of human cardiac ventricular action potential (Grandi et al., 2017; Sanguinetti et al., 1996). KCNQ1 also underlies K^+^ currents in the inner ear. Loss-of-function mutations in Q1 cause long QT syndrome type 1 (LQT1), exercise-induced sudden cardiac death, and deafness (Bohnen et al., 2017; Huang et al., 2018; Neyroud et al., 1997; Wang et al., 1996). While it is clear that Q1 stability and surface density are down-regulated by ubiquitination (Jespersen et al., 2007; Kanner et al., 2017), the role of distinct polyubiquitin linkage types in these processes are unknown.

Here, we developed a suite of engineered deubiquitinases (enDUBs) generated by fusing an anti-GFP/YFP nanobody to catalytic domains of DUBs with distinctive polyubiquitin chain preferences (OTUD1 -K63; OTUD4 - K48; Cezanne - K11; TRABID - K29/K33; USP21 - non-specific) (Mevissen et al., 2013; Mevissen and Komander, 2017; Ritorto et al., 2014), and applied these to investigate the roles of divergent polyubiquitin chains on KCNQ1-YFP stability, trafficking, and functional expression. A fraction of KCNQ1-YFP expressed in HEK293 cells was basally polyubiquitinated, with a dominance of K48 linkages followed by K63 and a lower level of other chain types (K11, K27, K33, K6) as assessed by mass spectrometry. Nevertheless, all the different enDUBs strongly decreased the total ubiquitin signal of KCNQ1-YFP. Despite the apparently similar impact on global KCNQ1-YFP ubiquitin content, the different enDUBs yielded sharply different effects on steady-state channel stability, surface density, subcellular localization, and rates of channel delivery to and removal from the surface membrane. Over-expressing either NEDD4-2 or ITCH with KCNQ1-YFP in HEK293 cells both resulted in a marked reduction of channel surface density and currents that was mediated through distinctive remodeling of the channel polyubiquitin linkage signature. Linkage-selective enDUBs had a different pattern of functional impact on Q1-YFP inhibited by either NEDD4-2 or ITCH compared to baseline, indicating context-dependent plasticity of the ubiquitin code. Expressing linkage-selective enDUBs with KCNQ1-YFP in cardiac myocytes yielded a unique pattern on channel stability and surface density that was different from observations in HEK293 cells, further emphasizing a cell-context-dependence of the ubiquitin code. Finally, individual trafficking-deficient LQT1 mutations in KCNQ1-YFP showed different patterns of rescue efficacy by linkage-selective enDUBs, revealing customized alterations of the ubiquitin code in response to substrate missense mutations. Overall, the results uncover a prominent role for distinct polyubiquitin chains in regulating KCNQ1 trafficking among separate subcellular compartments, demonstrate the plasticity of the ubiquitin code for a substrate depending on both extrinsic (cell context) and intrinsic (mutations) factors, and introduce the utility of linkage-selective enDUBs as a new tool to help demystify the polyubiquitin code.

## Results

### KCNQ1-YFP subcellular distribution and polyubiquitination in HEK293 cells

We used confocal microscopy and immunofluorescence to visualize steady-state distribution of KCNQ1-YFP in subcellular compartments – ER, Golgi, early endosomes (EE), late endosomes (LE), and lysosomes – after transient transfection in HEK293 cells. Most of the channel fluorescence co-localized with the ER, but we also observed significant co-localization with markers staining for the EE, LE, and lysosomal compartments (Fig. 1A,B). We observed a low degree of KCNQ1-YFP in the Golgi, potentially indicating a direct ER to plasma membrane trafficking mechanism (Nickel and Seedorf, 2008; Wilson et al., 2021), or a very short residence time in the Golgi (Fig. 1A,B). To selectively visualize and quantify plasma membrane channels, we engineered a 13-residue high-affinity bungarotoxin binding site (BBS) engineered into an extracellular loop of YFP-tagged KCNQ1 (BBS-KCNQ1-YFP) and incubated non-permeabilized transfected cells with Alexafluor-647-conjugated α-bungarotoxin (BTX-647) (Fig. 1C) (Aromolaran et al., 2014; Kanner et al., 2017). We used flow cytometry to rapidly measure YFP and BTX-647 fluorescence as metrics of total KCNQ1 expression and channel surface density, respectively (Fig. 1D). Transfected cells treated with MG132 to inhibit the proteasome moderately increased total and surface KCNQ1-YFP, enhanced channel co-localization with the ER and Golgi and decreased association with EE and lysosomal compartments (Fig. 1D; Supplemental Fig. 1). Notably, MG132 caused visually striking changes in the appearance of the ER (more clumped presentation) and lysosomes (Supplemental Fig. 1). By contrast with MG132, treating cells with chloroquine to eliminate lysosomal degradation yielded minimal effects on total and surface KCNQ1-YFP expression and subcellular localization (Supplemental Fig. 2). Thus, KCNQ1-YFP transiently expressed in HEK293 cells is differentially distributed among distinct subcellular compartments and degraded predominantly via the proteasome.

**Figure 1:**
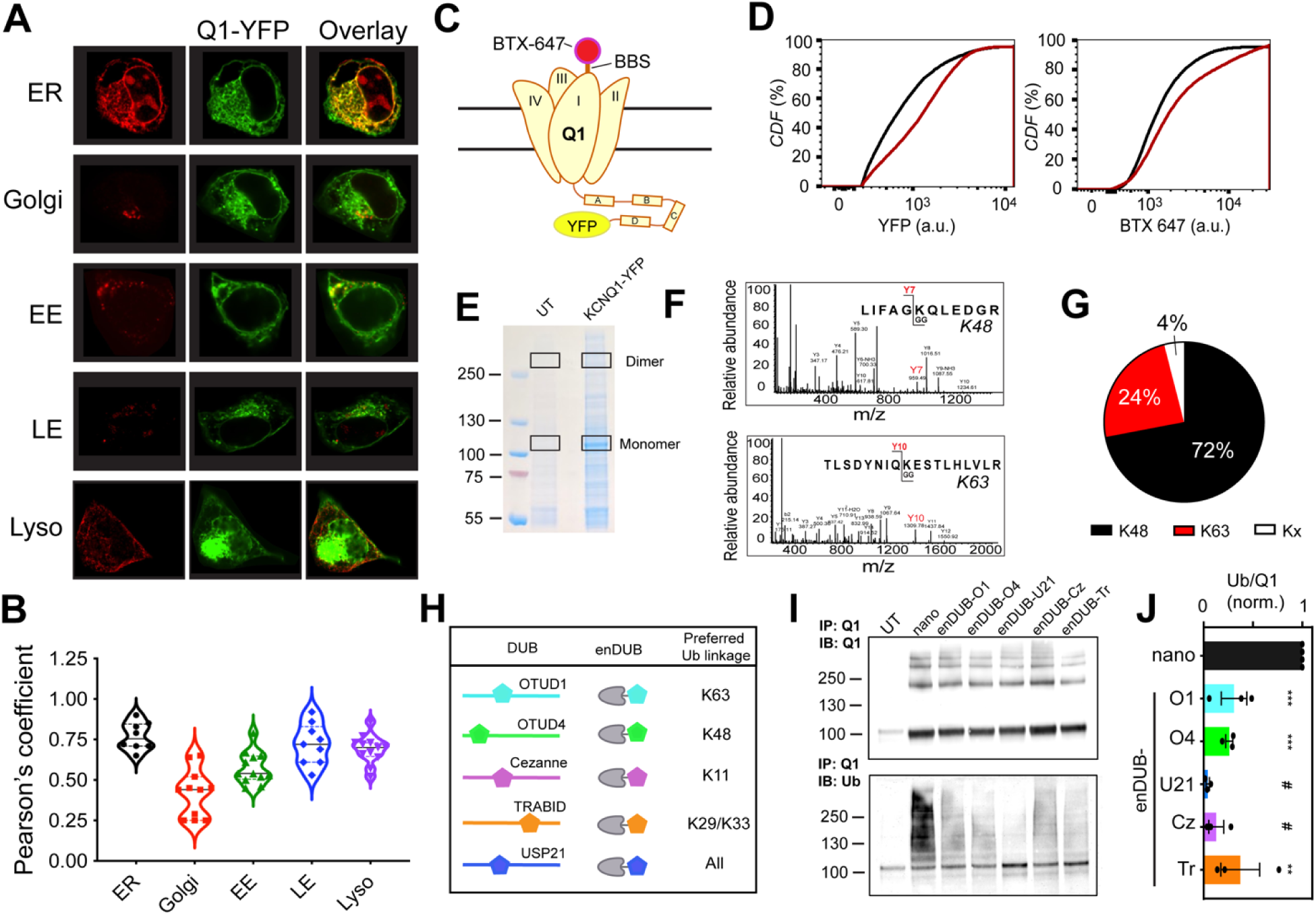
KCNQ1-YFP subcellular distribution and polyubiquitination in HEK293 cells and design of linkage-specific enDUBs. (A) Representative confocal images of HEK293 cells showing co-localization of expressed KCNQ1-YFP (green) and immunolabelled subcellular organelles [endoplasmic reticulum (ER), Golgi, early endosome (EE), late endosome (LE) and lysosome (lyso)] (red). (B) Co-localization of Q1-YFP with subcellular markers assessed by Pearson’s co-localization coefficient. (*N*=2 independent experiments; *n*>8 cells for each subcellular organelle imaged). (C) Schematic of BBS-KCNQ1-YFP with BTX-647 bound to an extracellular bungarotoxin-binding site (BBS). (D) Representative flow cytometry cumulative distribution function (CDF) plots showing total (YFP fluorescence) and surface (BTX-647) channels in cells expressing BBS-KCNQ1-YFP in untreated (black) and MG132 treated (red) conditions. (E) Representative Coomassie-stained SDS PAGE of pulled down KCNQ1-YFP. Black boxes indicate KCNQ1 monomeric and dimeric protein bands excised for mass spectrometry. (F) Representative ms2 spectra traces for ubiquitin peptides with di-glycine modification of K48 (*top*) and K63 (*bottom*), respectively. (G) Fractional distribution of Ub linkages on pulled down KCNQ1-YFP as assessed by mass spectrometry of associated di-glycine modified lysine residues on associated ubiquitin peptides. (H) Schematic of linkage-specific engineered deubiquitinases (enDUBs). (I) *Top*, KCNQ1-YFP pulldowns probed with anti-KCNQ1 in cells expressing KCNQ1-YFP with nano (control) or the indicated enDUBs. The four bands represent KCNQ1-YFP monomeric, dimeric, trimeric and tetrameric species. *Bottom*, same blot stripped and probed with anti-ubiquitin. (J) Relative KCNQ1-YFP ubiquitination computed as the ratio of anti-ubiquitin/anti-KCNQ1 signal intensities (mean ± SEM); *N*=3. One-way ANOVA and Dunnett’s multiple comparisons test, #*p*<0.0001, *** *p*<0.001 and ** *p*<0.01.

The precise molecular signals underlying KCNQ1 subcellular trafficking and degradation are unknown. We hypothesized that distinct polyubiquitin linkage chains could represent a prominent mechanism for controlling KCNQ1-YFP expression and trafficking among subcellular compartments. We used mass spectrometry to determine the prevalence of polyubiquitin chains on KCNQ1-YFP expressed in HEK293 cells. KCNQ1-YFP was pulled down in transiently transfected cells using anti-KCNQ1 antibody and detected by Coomassie staining following gel electrophoresis (Fig. 1E). Bands corresponding to KCNQ1-YFP monomer and dimers were excised for mass spectrometry analysis, using corresponding extracted segments from non-transfected cells as control (Fig. 1E). In addition to the expected KCNQ1 peptides, we identified enrichment of ubiquitin peptides compared to control, a fraction of which had di-glycines on lysine residues, a tell-tale signature of ubiquitin modification. Within the population of di-glycine modified ubiquitin lysine residues, K48 was fractionally dominant (72%) followed by K63 (24%), while other atypical chains (K11, K27, K29, K33, and K6) were much less prevalent (4%) (Fig. 1F,G; Supplemental Fig. 3).

### Rationale and design of linkage-selective and non-specific enDUBs

While the mass spectrometry results suggest the presence of distinct polyubiquitin linkages on KCNQ1-YFP expressed in HEK293 cells, this data does not provide any information on whether and how distinct polyubiquitin chain linkages regulate KCNQ1 expression, trafficking, and function. We sought to develop an approach to systematically examine the potential role of distinct polyubiquitin linkages in regulating KCNQ1 functional expression in live cells by exploiting minimal catalytic domains of deubiquitinases (DUBs) with preferences for hydrolyzing particular polyubiquitin linkages: OTUD1 (O1) - K63; OTUD4 (O4) - K48; Cezanne (Cz) - K11; TRABID (Tr) - K29/K33; and USP21 (U21) - non-specific (Mevissen et al., 2013; Mevissen and Komander, 2017; Ritorto et al., 2014). We fused the catalytic subunits of these DUBs to a GFP-targeted nanobody (Kubala et al., 2010) to generate putative linkage-selective engineered DUBs (enDUBs) that would act selectively on GFP/YFP-tagged protein substrates (Fig. 1H; Supplemental Fig. 4) (Kanner et al., 2020). We tested how effectively the different enDUBs decreased basal ubiquitination of KCNQ1-YFP. In control HEK293 cells expressing KCNQ1-YFP and the unmodified GFP nanobody (nano), immunoprecipitation followed by immunoblotting with anti-KCNQ1 yielded bands representing KCNQ1 monomers, dimers, trimers and tetramers (Fig. 1I, *top*). Stripping this blot and probing with anti-ubiquitin showed a smear consistent with basal KCNQ1 ubiquitination (Fig. 1I, *bottom*), and consistent with the mass spectrometry results (Fig. 1F,G). Co-expressing KCNQ1-YFP with the different enDUBs all resulted in significant decreases global in channel ubiquitination, albeit with quantitative differences in efficacy (Fig. 1I,J). In control experiments, catalytically dead enDUBs did not decrease KCNQ1-YFP ubiquitination, and in some cases appeared to increase ubiquitination, potentially reflecting a shielding effect (Supplemental Fig. 5).

### Differential impact of distinct enDUBs on KCNQ1 expression, subcellular localization, and function

The relatively uniform efficacy of all five enDUBs in reducing the amount of ubiquitin from KCNQ1-YFP (as reported by a pan-ubiquitin antibody) was somewhat surprising given the mass spectrometry data showing asymmetry in linkage types (Fig. 1G). There were three possibilities to explain this apparent discrepancy. First, in cells, the over-expressed enDUBs indiscriminately hydrolyzed all ubiquitin linkages in a substrate-independent manner. Second, prolonged tethering of the enDUBs to YFP enabled by the nanobody resulted in hydrolysis of all ubiquitin chain types present specifically on KCNQ1-YFP (Mevissen et al., 2013; Ritorto et al., 2014). Third, there is a prevalence of branched polyubiquitin chains with mixed linkages such that hydrolysis of even minor linkage species (atypical chains, such as K11, K29, K33, K6) can result in removal of a significant fraction of total ubiquitin associated with the channel. In the case of the first two possibilities, the distinct enDUBs would be expected to have a similar functional impact on channel expression and trafficking, whereas divergent effects could be possible in the third scenario. We assessed the impact of the different enDUBs on KCNQ1-YFP expression, trafficking, and function to distinguish among these possibilities.

We first used flow cytometry to contrast the impact of different enDUBs on BBS-KCNQ1-YFP total expression and surface density (Fig. 2A-E). Putatively targeting K63 hydrolysis with enDUB-O1 had no impact on either BBS-KCNQ1-YFP total expression or surface density. EnDUB-O4 (K48) also had no effect on KCNQ1 total expression, but unexpectedly reduced channel surface density. By contrast, enDUB-Cz (K11) and enDUB-Tr (K29/K33) both significantly increased KCNQ1 total expression and surface density (Fig. 2B-E). Finally, enDUB-U21 which hydrolyzes all chain types, increased KCNQ1 total expression without impacting channel surface density. In control experiments, none of the enDUBs affected BBS-KCNQ1 surface density when no YFP was attached to the channel, indicating the necessity of enDUB targeting to the substrate to yield a functional effect (Supplemental Fig. 6). The distinct enDUBs also produced characteristic changes to KCNQ1-YFP intracellular organelle distribution (Fig. 2F-J; Supplemental Fig. 7). By comparison to control (nano), enDUB-Cz and enDUB-Tr decreased co-localization with the ER, whereas enDUB-O4 increased ER retention (Fig. 2F); KCNQ1-YFP colocalization with the Golgi was strongly increased with enDUB-O4 and moderately elevated by enDUB-O1 and enDUB-U21 (Fig. 2G); enDUB-U21 decreased colocalization with LE (Fig. 2I); and enDUB-O1 diminished association with the lysosome (Fig. 2J).

**Figure 2:**
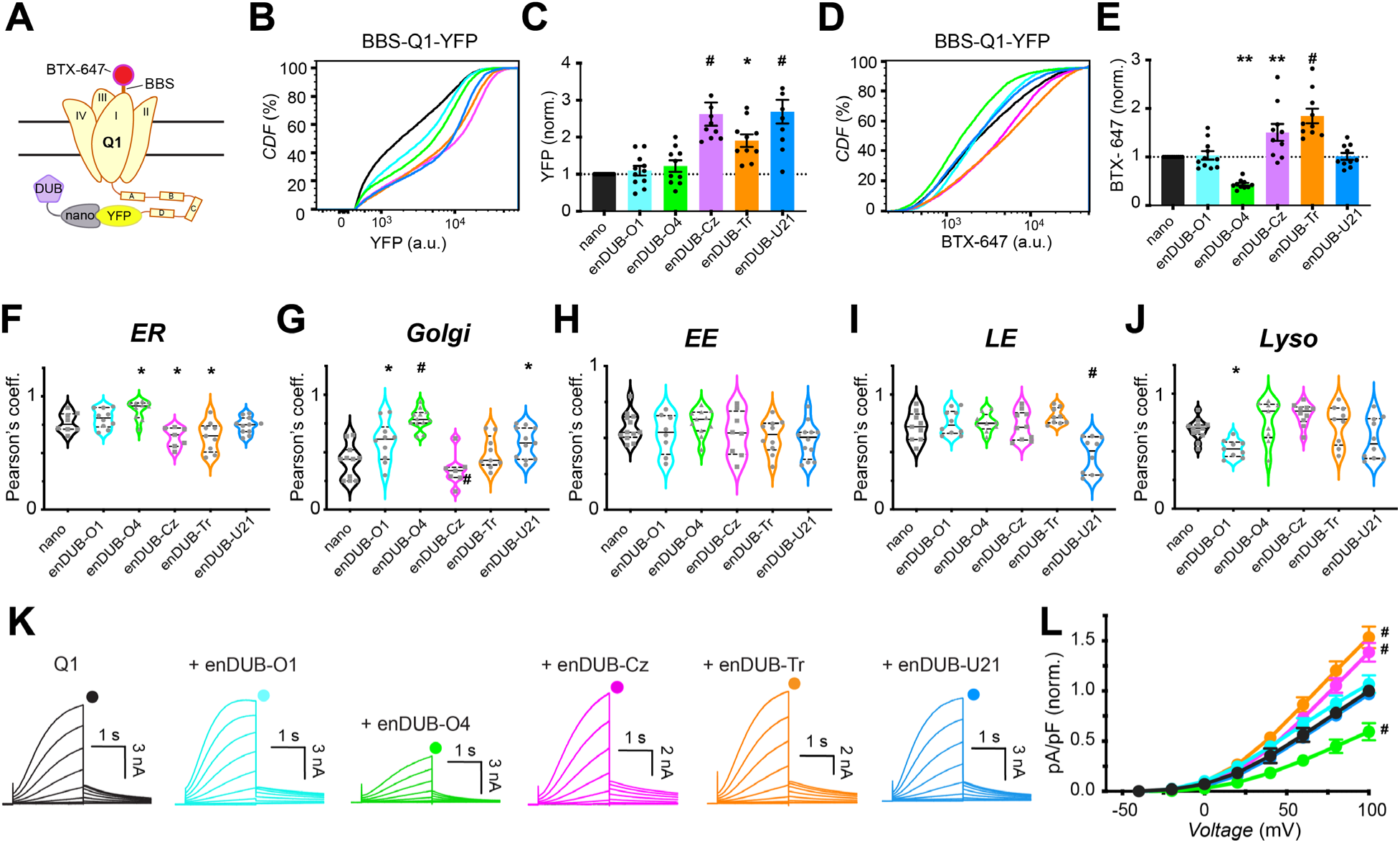
Differential impact of distinct enDUBs on KCNQ1 steady-state expression, subcellular localization, and function. (A) Experimental design schematic; BBS-KCNQ1-YFP is co-expressed with either GFP/YFP targeting nanobody (nano, control) or an enDUB in HEK293 cells. (B) Flow cytometry representative CDF plots showing total (YFP) fluorescence in cells expressing BBS-Q1-YFP and either nano alone (black) or each of the enDUBs (enDUB-O1, cyan; enDUB-O4, green; enDUB-Cz, magenta; enDUB-Tr, orange; enDUB-U21, blue). (C) Quantification of flow cytometry experiments for total KCNQ1-YFP expression analyzed from YFP-and CFP-positive cells (n>5,000 cells per experiment, *N*=10 independent experiments; #*p*<0.0001, \**p<0.05;* one-way ANOVA with Dunnett’s multiple comparisons). Data were normalized to values from the nano control group (dotted line). (D) Representative CDF plots showing surface fluorescence (BTX-647) in cells expressing BBS-KCNQ1-YFP and either nano alone (black) or an enDUB (enDUB-O1, cyane; enDUB-O4, green; enDUB-Cz, magenta; enDUB-Tr, orange; enDUB-U21, blue). (E) Quantification of flow cytometry experiments for KCNQ1-YFP surface expression analyzed from YFP-and CFP-positive cells (*n*>5,000 cells per experiment, *N*=10 independent experiments; #*p*<0.0001, \*\**p<0.01;* one-way ANOVA with Dunnett’s multiple comparisons). Data were normalized to values from the nano control group (dotted line). (F-J) Co-localization of KCNQ1-YFP with subcellular markers assessed by Pearson’s co-localization coefficient (*N*=2, *n*>8 for each subcellular organelle; #*p*<0.0001, \**p<0.05*, one-way ANOVA and Dunnett’s multiple comparisons test). (K) Exemplar KCNQ1 + KCNE1 current traces from whole-cell patch clamp measurements in CHO cells. (L) Population *I-V* curves for nano control (black, *n*=25), enDUB-O1 (cyan, *n*=9), enDUB-O4 (green, *n*=9), enDUB-Cz (magenta, *n*=16), enDUB-Tr (orange, *n*=16) and enDUB-U21 (blue, *n*=8). #*p*<0.0001, two-way ANOVA, with Tukey’s multiple comparisons. Here and throughout, data was pooled from *n* cells tested across three or more independent experimental days.

We next employed whole-cell patch clamp electrophysiology to determine the functional impact of the different enDUBs on KCNQ1 ionic currents (Fig. 2K,L). Control cells expressing KCNQ1-YFP + KCNE1 + nano displayed characteristic robust slowly activating delayed rectifier K^+^ currents (*I*_Ks_) (Fig. 2K) (Aromolaran et al., 2014; Sanguinetti et al., 1996). In concordance with the observed effects on channel surface density, enDUB-O1 and enDUB-U21 had minimal effects on *I*_Ks_, enDUB-O4 significantly decreased *I*_Ks_, while enDUB-Cz and enDUB-Tr both increased *I*_Ks_ (Fig. 2K,L).

Overall, these results revealed a number of unexpected findings including that: hydrolysis of apparently minor polyubiquitin linkage chains species (K11 and K29/K33) yielded the most robust stabilization of KCNQ1 expression; hydrolysis of different polyubiquitin linkages distinctively alter KCNQ1 distribution among various intracellular compartments; that hydrolysis of distinct ubiquitin chains can selectively suppress or upregulate KCNQ1 surface density independent of effects on channel expression; and that K48 polyubiquitin, the well-established canonical degradation signal, does not control KCNQ1 expression in this context but rather is important for channel trafficking to the cell surface. The results also validate the linkage-selective enDUB strategy as capable of providing unique insights into the *in situ* functions of distinct polyubiquitin linkage types in a substrate-specific manner.

### Distinctive impact of enDUBs on KCNQ1 delivery to and removal from the plasma membrane

The steady-state surface density of KCNQ1 is maintained by dynamic processes of channel delivery to and removal from the plasma membrane. KCNQ1 delivery to the surface involves both the transport of newly synthesized channels via the Golgi (or directly from the ER) and the recycling of a fraction of endocytosed channels. We wondered whether the divergent effects of the enDUBs on KCNQ1 steady-state surface density was mediated by differential regulation of dynamic forward trafficking and internalization processes. To address this question, we utilized optical-pulse chase assays to determine rates of BBS-KCNQ1-YFP delivery to and removal from the plasma membrane and assessed the impact of the enDUBs on these processes (Fig. 3).

**Figure 3:**
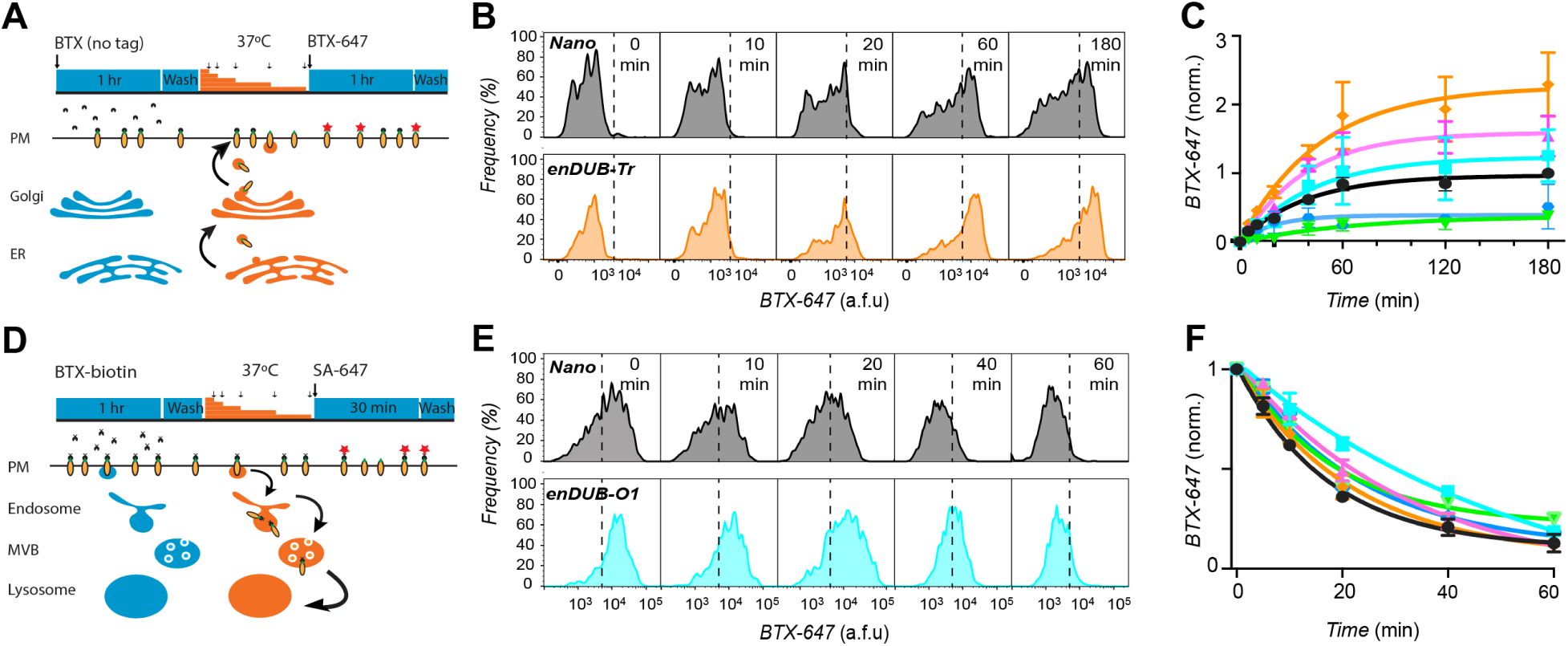
Distinctive impact of different enDUBs on dynamic KCNQ1 delivery to and removal from the plasma membrane. (A) Schematic of optical pulse-chase assay to measure BBS-KCNQ1-YFP forward trafficking. (B) Representative flow cytometry BTX-647 fluorescence histograms showing time evolution of surface channel increase in cells expressing BBS-KCNQ1-YFP with nano (black, *top*) or enDUB-Tr (orange, *bottom*). (C) Time evolution of channel forward to the surface in cells expressing BBS-KCNQ1-YFP with either nano or an enDUB (enDUB-O1, cyan; enDUB-O4, green; enDUB-Cez, magenta; enDUB-Tr, orange; enDUB-U21, blue; *n*>10,000 cells; *N*=3 for each data point; *p*>0.05, nano vs enDUB-O1; *p*<0.0001, nano vs enDUB-O4; *p*<0.01, nano vs enDUB-Cz; *p*<0.0001, nano vs enDUB-Tr; *p*<0.01, nano vs enDUB-U21; two-way ANOVA, with Tukey’s multiple comparisons). Data are normalized to the max value of nano control (black). Smooth curves are fits of an exponential growth function to the data: y= Ae^1/τ^ +y0. (D) Schematic of endocytosis pulse-chase assay. (E) Representative flow cytometry BTX-647 fluorescence histograms showing time evolution of surface channel decrease in cells expressing BBS-KCNQ1-YFP with nano (black, *top*) or enDUB-O1 (turquoise, bottom). (F) Time evolution of channel removal from the surface in cells expressing BBS-KCNQ1-YFP with either nano or an enDUB (enDUB-O1, cyan; enDUB-O4, green; enDUB-Cz, magenta; enDUB-Tr, orange; enDUB-U21, blue; *n*>10,000 cells; *N*=3 for each data point; *p*<0.0001, nano vs enDUB-O1, *p*<0.05, nano vs enDUB-O4, *p*<0.05, nano vs enDUB-Cz, *p*>0.05, nano vs enDUB-Tr, *p*>0.05, nano vs enDUB-U21, two-way ANOVA with Tukey’s multiple comparisons). Smooth curves are fits of an exponential growth function to the data: y= Ae^-1/τ^ +y0.

To monitor channel delivery to the plasma membrane, HEK293 cells transiently expressing BBS-Q1-YFP were initially incubated at 4°C to arrest trafficking processes after which the non-permeabilized cells were exposed to excess unconjugated (non-fluorescent) BTX to saturate extracellular BBS epitopes (pulse). Cells were then incubated at 37°C for varying times to resume trafficking (chase), and the newly delivered BBS-Q1-YFP channels labeled with BTX-647 at 4°C, followed by quantifying fluorescence signals using flow cytometry (Fig. 3A) (Kanner et al., 2017). In control cells expressing BBS-Q1-YFP, the BTX-647 increased exponentially with a half-time of 20.37 mins, and reaching a plateau after 2 hrs (Fig. 3B,C). To enable facile comparison among different experiments, we normalized BTX-647 signals to the maximum value obtained from the contemporaneous control group (at 3 hrs), following background subtraction (Fig. 3C). The dominant effects of the distinct enDUBs lay in their dramatically different effects on the amount of BBS-KCNQ1-YFP delivered to the cell surface, as reported by relative plateau levels of the forward trafficking curves (Fig. 3C). Whereas enDUB-O1 had only a marginal effect, enDUB-O4 and enDUB-U21 both led to a marked ∼70% decrease in the BTX-647 steady-state signal compared to control, reflecting a strong suppression of channel forward trafficking. In sharp contrast, enDUB-Cz and enDUB-Tr both robustly enhanced KCNQ1 forward trafficking as revealed by 1.6 and 2-fold increases, respectively, in BTX-647 plateau signal relative to control (Fig. 3C).

To measure KCNQ1 endocytosis, BBS-KCNQ1-YFP channels initially at the cell surface were labeled with biotinylated bungarotoxin (BTX-biotin) at 4°C (pulse). Cells were then incubated at 37°C for varying time periods to resume trafficking (chase), followed by labeling with streptavidin-conjugated Alexa Fluor 647 (SA-647) at 4°C (Fig. 3D) (Kanner et al., 2017). In this paradigm, only channels that were present at the cell surface during the pulse phase and labeled with BTX-biotin would be fluorescently labeled with SA-647 after the chase, distinguishing them from new channels that are subsequently inserted. A decrease in SA-647 fluorescence signal with increasing chase times would be expected due to internalization of BTX-biotin-labeled channels. Indeed, control cells expressing BBS-KCNQ1-YFP + nano displayed an exponential decline in SA-647 fluorescence with chase time, with a half-life (*t*_1/2_) of 12.02 mins (Fig. 3E, F). The different enDUBs produced distinctive influences on the apparent rate of BBS-KCNQ1-YFP internalization (Fig. 3F). The most dramatic effect was observed with enDUB-O1 which slowed the apparent rate of internalization by over 3-fold (*t*_1/2_ = 38.32 mins) (Fig. 3E,F). EnDUB-Cz caused an intermediate slowing of the apparent rate of internalization (*t*_1/2_ = 21.79 mins), whereas enDUB-O4 (*t*_1/2_ = 12.43 mins), enDUB-Tr (*t*_1/2_ = 15.06 mins), and enDUB-U21 (*t*_1/2_ = 15.61 mins) had marginal effects (Fig. 3F).

Recycling of KCNQ1-YFP is another layer of subcellular dynamics that may be modulated by ubiquitin and may potentially contribute to the measurements of forward trafficking and endocytosis. To determine the potential contribution of channel recycling to BBS-KCNQ1-YFP steady-state surface density, we utilized dominant negative Rab11 (Rab11-DN) to block slow recycling endosomes (Supplemental Fig. 8). We observed a ∼20% decrease in BBS-KCNQ1-YFP steady-state density in cells co-expressing Rab11-DN compared to nano (control) (Supplemental Fig. 8). In the presence of Rab11-DN, enDUB-O1 yielded a 50% suppression in BBS-KCNQ1-YFP surface density (Supplemental Fig 8). Co-expressing Rab11-DN also largely neutralized the increase in surface density observed with enDUB-Cz (Supplemental Fig. 8). These data indicate that under baseline conditions (i.e. without Rab11-DN) enDUB-O1 and enDUB-Cz increase the pool of channels recycled to the cell surface after endocytosis. Thus, by inference, K63 and K11 reduce the pool of channels recycled to the surface. By contrast, Rab11-DN did not substantively diminish the increase in BBS-KCNQ1-YFP observed with enDUB-Tr (Fig. 2D,E; Supplemental Fig. 8) suggesting a minimal impact of K29/K33 on channel recycling. Altogether, these data suggest that pruning K11 enhances forward trafficking from ER/Golgi, slows endocytosis, and promotes recycling; removing K29/K33 from the channel solely enhances forward trafficking from the ER/Golgi; whereas stripping K63 slows endocytosis and increases channel recycling.

### KCNQ1 is downregulated by NEDD4L and ITCH with distinct polyubiquitin signatures

The system of writers, erasers, readers, and editors that constitute the ubiquitin code undoubtedly varies among different cell types due to expected differences in protein expression profiles. As such, the ubiquitin code regulation of a particular protein may not be immutable but rather change in different cellular environments. To explore this anticipated but under-explored dimension of the ubiquitin code, and to further test the discriminative capacity of linkage-selective enDUBs, we sought to perturb the polyubiquitin signature of KCNQ1 in HEK293 cells. To this end, the HECT E3 ligase NEDD4-2 which predominantly catalyzes the synthesis of K63 polyubiquitin chains on substrates, is a known regulator of KCNQ1 (Jespersen et al., 2007; Kim and Huibregtse, 2009). Accordingly, NEDD4-2 co-expression dramatically reduced both KCNQ1 surface density (Fig. 4A,B) and whole-cell currents (Fig. 4C,D). We screened other members of the NEDD4 family of HECT E3 ligases and identified ITCH as a novel regulator of KCNQ1. Similar to NEDD4-2, ITCH significantly down-regulated both KCNQ1 channel surface density and ionic currents (Fig. 4A-D). The functional effects of NEDD4-2 were diminished by a mutation within a canonical PY motif located on the KCNQ1 C-terminus (Y662A) (Supplemental Fig. 9) as previously reported (Jespersen et al., 2007). Surprisingly, the functional effects of ITCH on KCNQ1 were unaffected by the Y662A mutation (Supplemental Fig. 9), indicating a potential difference in how it associates with the channel compared to NEDD4-2.

**Figure 4:**
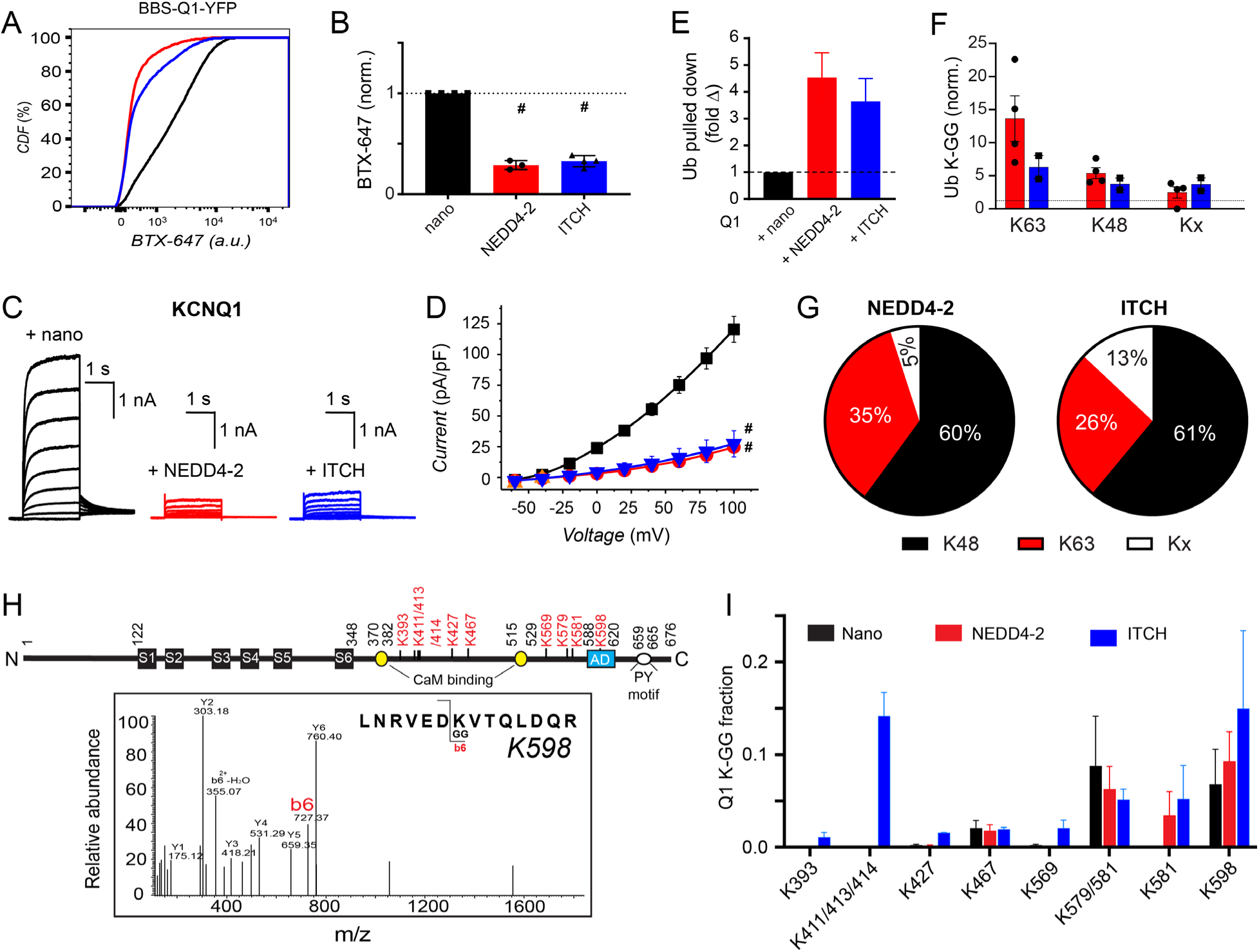
KCNQ1 functional expression is downregulated by the E3 ligases NEDD4-2 and ITCH with distinct polyubiquitin signatures. (A) Representative flow cytometry CDF plots showing surface channel (BTX-647) fluorescence in cells expressing BBS-KCNQ1-YFP and nano (black) with NEDD4L (red) or ITCH (blue). (B) Mean BBS-KCNQ1-YFP surface expression (BTX-647 fluorescence) analyzed from YFP-positive cells (*n*>5,000 cells per experiment, *N*=3-4; *#p*<0.0001, one-way ANOVA and Dunnett’s multiple comparisons test). Data were normalized to values from the nano control group (dotted line). (C) Exemplar KCNQ1 current traces from whole-cell patch clamp measurements in CHO cells. (D) Population *I-V* curves for KCNQ1 + KCNE1 + nano (control; black, *n*=10), KCNQ1+ NEDD4-2 (red, *n*=9), and KCNQ1+ITCH (blue, *n*=11) (#*p*<0.0001, two-way ANOVA, with Tukey’s multiple comparisons). (E) Quantification of ubiquitin that was pulled down with KCNQ1-YFP expressed with nano (control), NEDD4-2 or ITCH, as assessed by mass spectrometry. Data were normalized to the control group. (F) Mass spectrometric evaluation of ubiquitin lysine residues that are di-gly modified (*n*=4, *N*=2). (G) Fractional distribution of Ub linkages as assessed by mass spectrometric analysis of immunoprecipitated and gel excised bands of KCNQ1-YFP co-expressed with NEDD4L or ITCH. (H) Cartoon of KCNQ1 with di-gly-modified lysine residues identified by mass spectrometry shown in red. Representative ms2 spectra trace of KCNQ1 peptide with di-gly modification of K598. (I) Quantification of di-gly-modified lysine residues on KCNQ1 peptides (*N*=2 independent experiments).

We used mass spectrometry to determine the impact of NEDD4-2 and ITCH co-expression, respectively, on KCNQ1 polyubiquitin signatures. Mass spectrometry analyses on KCNQ1 pulldown bands excised from a Coomassie stained gel indicated that NEDD4-2 and ITCH co-expression, respectively, yielded 4.5-fold and 3.5-fold increases in total ubiquitin associated with the channel compared to control (nano) (Fig. 4E). Compared to control, NEDD4-2 increased the relative amount of ubiquitin peptides with di-glycine modifications on K63 (13-fold), K48 (5-fold), and other linkages, Kx (3-fold) (Fig. 4F). ITCH also increased di-glycine modifications of ubiquitin K63 (6-fold), K48 (4-fold), and Kx (4-fold) (Fig. 4F). Further, in the ubiquitin pool pulled down with the channel, NEDD4L significantly increased the fraction of K63 ubiquitin linkages (from 24% to 35%) and decreased K48 (from 72% to 60%) while having a modest effect on the fraction of other linkage types (from 4% to 5%) (Fig. 4G) relative to control (Fig. 1G). By contrast, ITCH significantly increased the fraction of atypical linkages Kx (from 4% to 13%), marginally altered the fraction of K63 (from 24% to 26%), and reduced K48 (from 72% to 61%) (Fig. 4G) relative to control (Fig. 1G).

Beyond variations in polyubiquitin linkage types, another potential factor influencing the functional impact of NEDD4-2 and ITCH on KCNQ1-YFP functional expression is the complement of lysine residues on the channel that are ubiquitinated under the different conditions. There are 31 intracellular lysine residues on KCNQ1 that could potentially act as sites for ubiquitin attachment of which 25 are present in the C-terminus (Supplemental Fig. 10). In control cells expressing KCNQ1-YFP + nano, mass spectrometry indicated three lysine residues – K467, K579, and K598 – in which the fraction of di-glycine modified peptide exceeded 1% (Fig. 4H,I). NEDD4-2 did not appreciably change the fraction of di-glycine modified K467, K579, and K598, but did result in the emergence of di-glycine modified K581 (Fig. 4H,I). By contrast, ITCH yielded detection of more di-glycine modified lysines on KCNQ1 including K393, K411/K412/K413, K427 and K569 in addition to K467, K579, K581 and K598 seen with NEDD4-2 (Fig. 4H,I).

Overall, these data suggest that the similar functional effects of NEDD4-2 and ITCH on KCNQ1 currents and surface density may be mediated through divergent polyubiquitin signatures. Thus, co-expression of NEDD4-2 and ITCH constitute unique adjustments of the ubiquitin code regulating KCNQ1 functional expression, providing a forum to further test the discriminatory capacity of linkage-selective enDUBs.

### Differential impact of enDUBs on reversing KCNQ1 functional downregulation by NEDD4L and ITCH

We first investigated the capacity of the different enDUBs to counter the impact of NEDD4-2 on KCNQ1 ubiquitination and functional expression (Fig 5). In control cells co-expressing KCNQ1-YFP + NEDD4-2 + nano, pulldown of the channel followed by immunoblotting with KCNQ1 antibodies yielded specific bands consistent with full-length monomeric KCNQ1-YFP and oligomeric species (Fig. 5B, *left*). Stripping the blot and probing with ubiquitin antibodies yielded strong staining (Fig. 5B, *right*), consistent with the ∼4.5-fold increase in KCNQ1-YFP ubiquitination in the presence of NEDD4-2 overexpression observed by mass spectrometry (Fig. 4E). All five enDUBs when coexpressed with KCNQ1-YFP + NEDD4-2 dramatically reduced the ubiquitin signal on pulled-down KCNQ1-YFP (Fig. 5B,C). Despite the apparently similar impact of the distinct enDUBs on the broad biochemical signature of ubiquitin pulled down with the channel, analyses of their effects on channel expression, subcellular localization, and ionic currents delineated sharp differences amongst them. KCNQ1 expression was strongly enhanced by enDUB-Cz/enDUB-U21, more moderately increased by enDUB-O1/enDUB-Tr, and decreased by enDUB-O4 (Fig. 5D,E). The decrease in KCNQ1 surface density caused by NEDD4-2 overexpression was reversed by enDUB-O1 and enDUB-Cz to beyond the level observed under control baseline conditions (Fig. 5F,G). Channel surface density was also rescued by enDUB-U21, although to a smaller extent than observed with enDUB-O1/enDUB-Cz. Surprisingly, enDUB-Tr was completely ineffective in rescuing surface density of KCNQ1-YFP channels down-regulated by NEDD4-2 overexpression (Fig. 5F,G). Whole-cell patch clamp studies further consolidated these results. Consistent with the pattern of rescue of KCNQ1 surface density, *I*_Ks_ currents suppressed by NEDD4-2 over-expression were strongly rescued by enDUB-O1/enDUB-Cz/enDUB-U21 and refractory to enDUB-O4 and enDUB-Tr (Fig. 5H,I).

**Figure 5:**
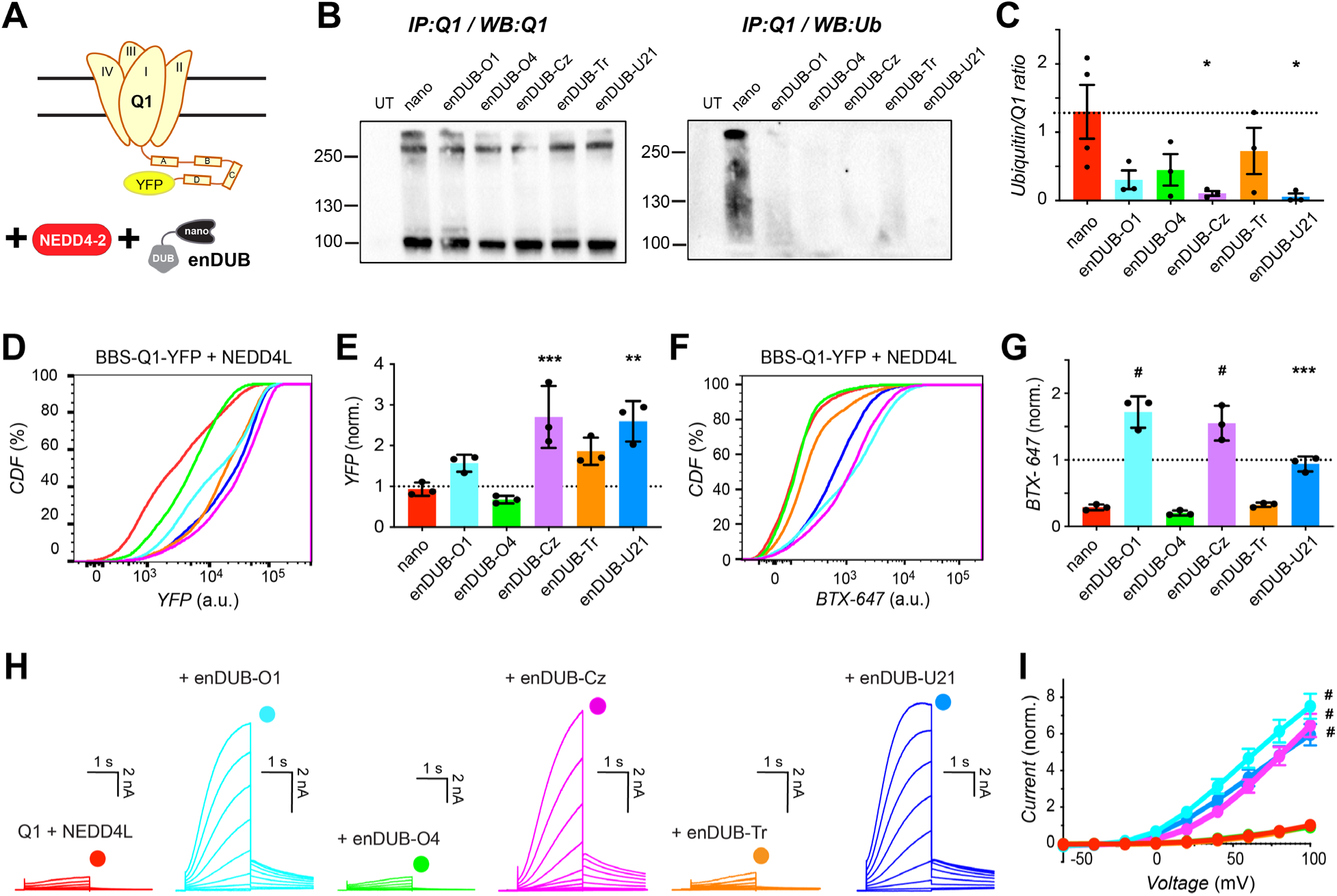
Differential impact of distinct linkage-specific enDUBs on reversing KCNQ1 functional downregulation by NEDD4-2. (A) Experimental design schematic; BBS-KCNQ1-YFP is co-expressed with NEDD4-2 and either nano (control) or an enDUB in HEK293 cells. (B) *Left*, Western blot of KCNQ1-YFP pulldowns from cells expressing KCNQ1-YFP + NEDD4-2 and either nano or the indicated enDUB, probed with anti-KCNQ1 antibody. *Right*, same blot stripped and probed with anti-ubiquitin. (C) Relative KCNQ1 ubiquitination computed by the ratio of anti-ubiquitin/anti-KCNQ1 signal intensities; (*n*=3); \**p*<0.05, one-way ANOVA with Dunnett’s multiple comparisons. (D) Flow cytometry representative CDF plots showing total channel expression (YFP fluorescence) in cells expressing BBS-KCNQ1-YFP + NEDD4-2 and either nano or an enDUB. (E) Mean KCNQ1 expression analyzed from YFP-and CFP-positive cells. Data were normalized to values from the nano control group (dotted line) (*n*>5,000 cells per experiment; *N*=3; \*\*\**p*<0.001 and \*\**p*<0.01, one-way ANOVA with Dunnett’s multiple comparisons). (F) Representative CDF plots showing surface channels (BTX-647 fluorescence) in cells expressing BBS-KCNQ1-YFP + NEDD4-2 and either nano or an enDUB. (G) Mean BBS-KCNQ1-YFP surface expression analyzed from YFP-and CFP-positive cells. Data were normalized to values from the nano control group (dotted line) (*n*>5,000 cells per experiment; *N*=3; \*\*\**p*<0.001 and #*p*<0.0001, one-way ANOVA with Dunnett’s multiple comparisons). (H) Exemplar KCNQ1 + KCNE1 currents from whole-cell patch clamp measurements in CHO cells. (I) Population *I-V* curves for nano (control; red, *n*=18), enDUB-O1 (turquoise, *n*=9), enDUB-O4 (green, *n*=9), enDUB-Cz (pink, *n*=10), enDUB-Tr (orange, *n*=10) and enDUB-U21 (blue, *n*=8). #*p*<0.0001, two-way ANOVA, with Tukey’s multiple comparisons.

In cells co-expressing KCNQ1-YFP + ITCH, all five enDUBs again effectively decreased the amount of ubiquitin pulled-down with the channel (Fig. 6A-C). KCNQ1-YFP expression in the presence of ITCH was increased by enDUB-Cz, enDUB-Tr, and enDUB-U21, while enDUB-O1 and enDUB-O4 had minimal impact (Fig. 6D,E). KCNQ1 surface density and whole-cell currents suppressed by ITCH was strongly increased by enDUB-O1, enDUB-Cz, and enDUB-U21, while enDUB-Tr yielded an intermediate rescue (Fig. 6G-I). The partial effect of enDUB-Tr in KCNQ1 channels downregulated by ITCH contrasted sharply with the lack of an effect observed with NEDD4-2.

**Figure 6:**
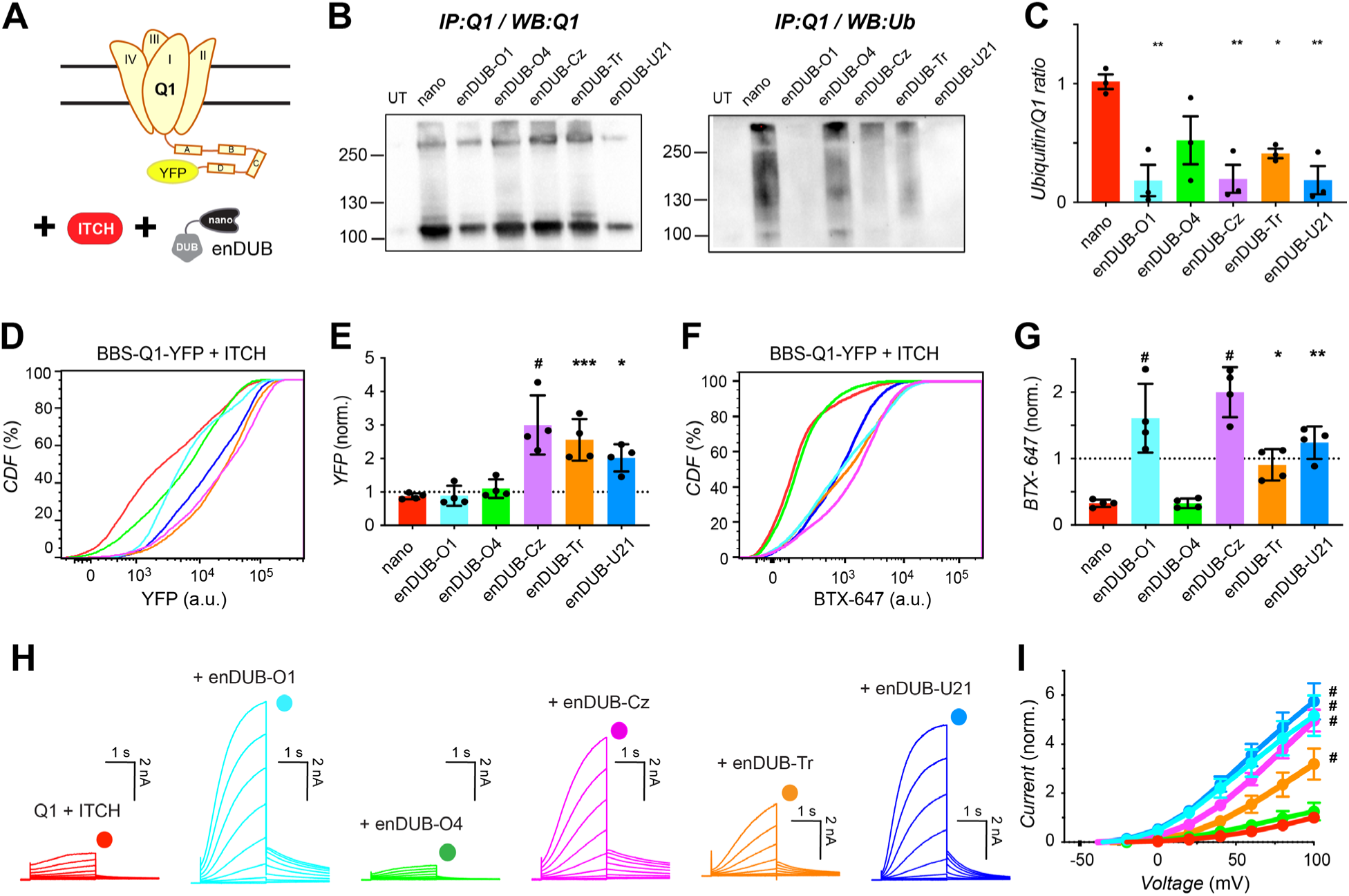
Differential impact of distinct linkage-specific enDUBs on reversing KCNQ1 functional downregulation by ITCH. (A) Experimental design schematic; BBS-KCNQ1-YFP is co-expressed with ITCH and either nano (control) or an enDUB in HEK293 cells. (B) *Left*, Western blot of KCNQ1-YFP pulldowns from cells expressing KCNQ1-YFP + ITCH and either nano or the indicated enDUB, probed with anti-KCNQ1 antibody. *Right*, same blot stripped and probed with anti-ubiquitin. (C) Relative KCNQ1 ubiquitination computed by the ratio of anti-ubiquitin/anti-KCNQ1 signal intensities (*n*=3; \*\**p*<0.01 and \**p*<0.05, one-way ANOVA with Dunnett’s multiple comparisons). (D) Flow cytometry representative CDF plots showing total channel expression (YFP fluorescence) in cells expressing BBS-KCNQ1-YFP + ITCH and either nano or an enDUB. (E) Mean KCNQ1 expression analyzed from YFP-and CFP-positive cells (*n*>5,000 cells per experiment; *N*=3; #*p*<0.0001, \*\*\**p*<0.001 and \**p*<0.05, one-way ANOVA with Dunnett’s multiple comparisons). Data were normalized to values from the nano control group (dotted line). (F) Representative CDF plots showing surface channels (BTX-647 fluorescence) in cells expressing BBS-KCNQ1-YFP + NEDD4-2 and either nano or an enDUB. (G) Mean BBS-KCNQ1-YFP surface expression analyzed from YFP-and CFP-positive cells (*n*>5,000 cells per experiments; *N*=3; #*p*<0.0001, \*\**p*<0.01 and \**p*<0.05, one-way ANOVA with Dunnett’s multiple comparisons). Data were normalized to values from the nano control group (dotted line). (H) Exemplar KCNQ1 + KCNE1 currents from whole-cell patch clamp measurements in CHO cells. (I) Population *I-V* curves for nano (control; red, *n*=18), enDUB-O1 (turquoise, *n*=9), enDUB-O4 (green, *n*=8), enDUB-Cez (pink, *n*=10), enDUB-Tr (orange, *n*=10) and enDUB-U21 (blue, *n*=8). #*p*< 0.0001, two-way ANOVA with Tukey’s multiple comparisons.

Overall, the pattern of effects of distinct enDUBs on KCNQ1 surface density and currents differs between the basal, NEDD4-2 and ITCH overexpression conditions, emphasizing the adjustability of the ubiquitin code in response to changes in the cellular context, and further affirming the utility of linkage-selective enDUBs as tools to decipher the polyubiquitin code regulation of protein substrates *in situ*.

### Differential impact of linkage-selective enDUBs on KCNQ1 functional expression in cardiomyocytes

The malleability of the polyubiquitin code regulation of KCNQ1 revealed by our experiments suggested that polyubiquitin linkage regulation of the channel could potentially vary substantively in different cell types. As such, it was of interest to probe the putative role of distinct polyubiquitin chain linkages on functional expression of KCNQ1 within the native context of ventricular cardiomyocytes. We generated adenoviral vectors expressing BBS-KCNQ1-YFP and the different enDUBs in a bicistronic IRES-mCherry format, and used these to infect acutely cultured adult guinea pig ventricular cardiomyocytes. We utilized confocal microscopy to visualize total channel expression (YFP fluorescence), nano or enDUB expression (mCherry fluorescence), and surface channels (Alexa fluor 647 fluorescence) (Fig. 7A).

**Figure 7:**
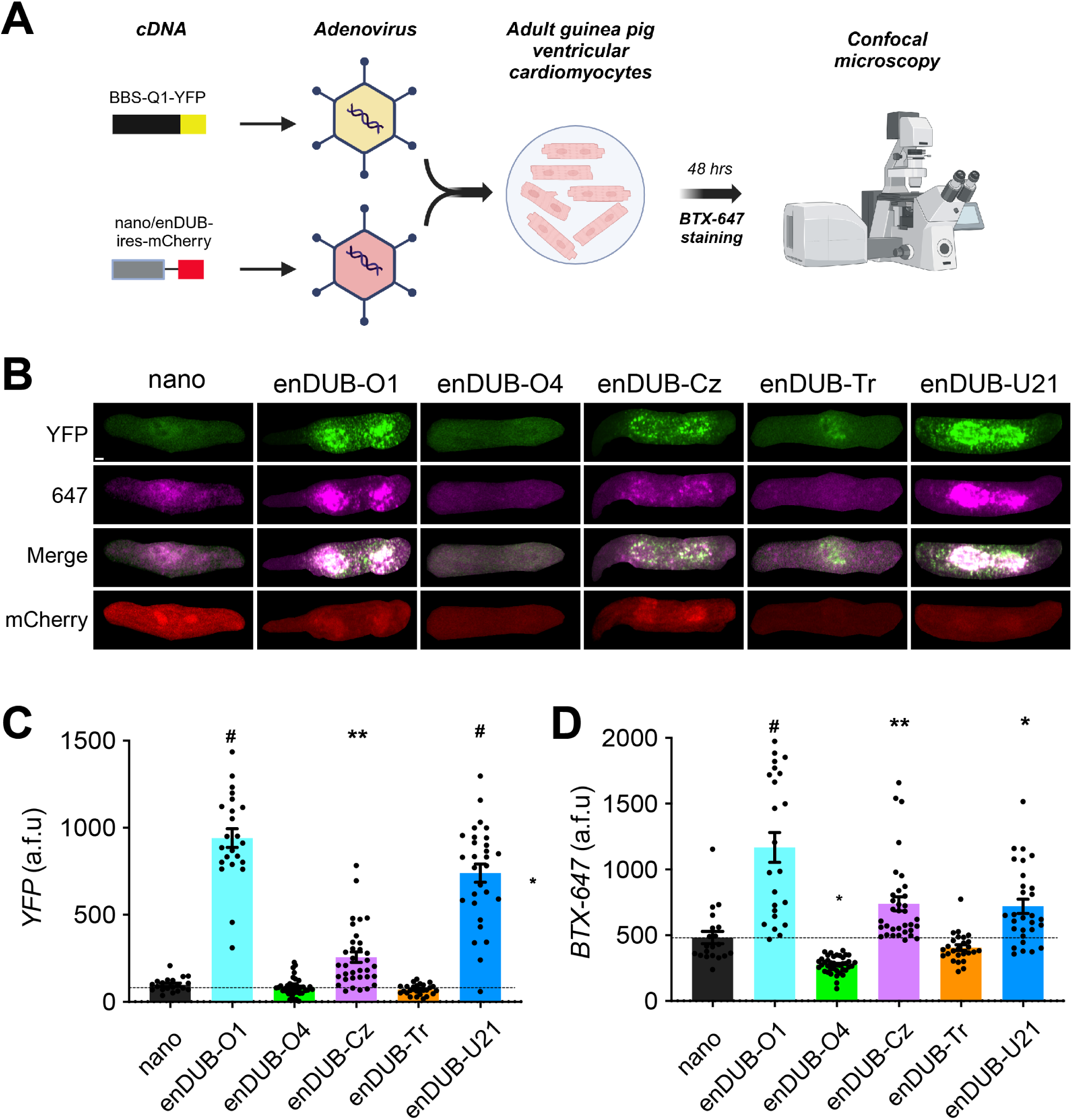
Differential impact of linkage-specific enDUBs on KCNQ1 functional expression in cardiomyocytes. (A) Schematic of the experimental design. (B) Representative confocal images of adult guinea pig ventricular cardiomyocytes expressing BBS-KCNQ1-YFP with either nano-IRES-mCherry or an enDUB-IRES-mCherry, showing YFP (top row), BTX-647 (second row), YFP and BTX-647 overlay (third row), and mCherry (bottom) fluorescence signals. Scale bar, 20 µm. (C) Quantification of YFP fluorescence intensity (total channel expression) measured from confocal images (*N*=3 isolations; mean and SEM; *#p*< 0.0001 and **p<0.01, one-way ANOVA and Dunnett’s multiple comparisons test). (D) Quantification of the surface BTX-647 fluorescence intensity (surface channels) measured from confocal images (*N*=3 isolations; mean and SEM; *#p*<0.0001, *p<0.05 and **p<0.01, one-way ANOVA and Dunnett’s multiple comparisons test).

Control cells expressing BBS-Q1-YFP + nano-IRES-mCherry displayed robust YFP, mCherry, and 647 fluorescence signals indicating a fraction of the channels were present at the cardiomyocyte surface (Fig. 6B-D). Intriguingly, enDUB-O1 and enDUB-U21 strongly boosted KCNQ1 expression (YFP fluorescence), enDUB-Cz had an intermediate effect, while enDUB-O4 and enDUB-Tr had no impact (Fig. 6B,C). Channel surface density was strongly enhanced by enDUB-O1 while enDUB-Cz and enDUB-U21 had intermediate effects. By contrast, enDUB-O4 decreased surface density while enDUB-Tr had no impact (Fig. 6B,D).

Thus, the signature pattern of distinct enDUBs effects on KCNQ1 expression and surface density in cardiomyocytes differs qualitatively and quantitatively from observations in HEK293 cells, further demonstrating the cell-context-dependent plasticity of the ubiquitin code.

### KCNQ1 disease mutants alter signature pattern of responsiveness to distinct enDUBs

Loss-of-function mutations in KCNQ1 that perturb channel trafficking cause LQT1 and may also lead to deafness. We previously showed that targeted deubiquitination with an enDUB can rescue channel surface density and ionic currents in a subset of trafficking-deficient KCNQ1 mutations (Kanner et al., 2020). However, it is unknown whether such KCNQ1 missense mutations alter polyubiquitin linkage regulation of the channel. Accordingly, we tested the impact of the different enDUBs on expression and surface density of four individual LQT1 mutations localized to the KCNQ1 C terminus— R366W, R539W, T587M and G589D (Fig. 8A). The mutations mostly had relatively modest impact on KCNQ1 total expression, with only T587M yielding a 30% decrease in YFP fluorescence intensity (Fig. 8F). The signature pattern of enDUB effects on expression levels of the mutants was similar to wild type KCNQ1 under basal conditions— enDUB-Cz, enDUB-Tr, and enDUB-U21 all increased YFP fluorescence whereas enDUB-O1 and enDUB-O4 had minimal effects (Fig. 8B,D,F,H).

**Figure 8:**
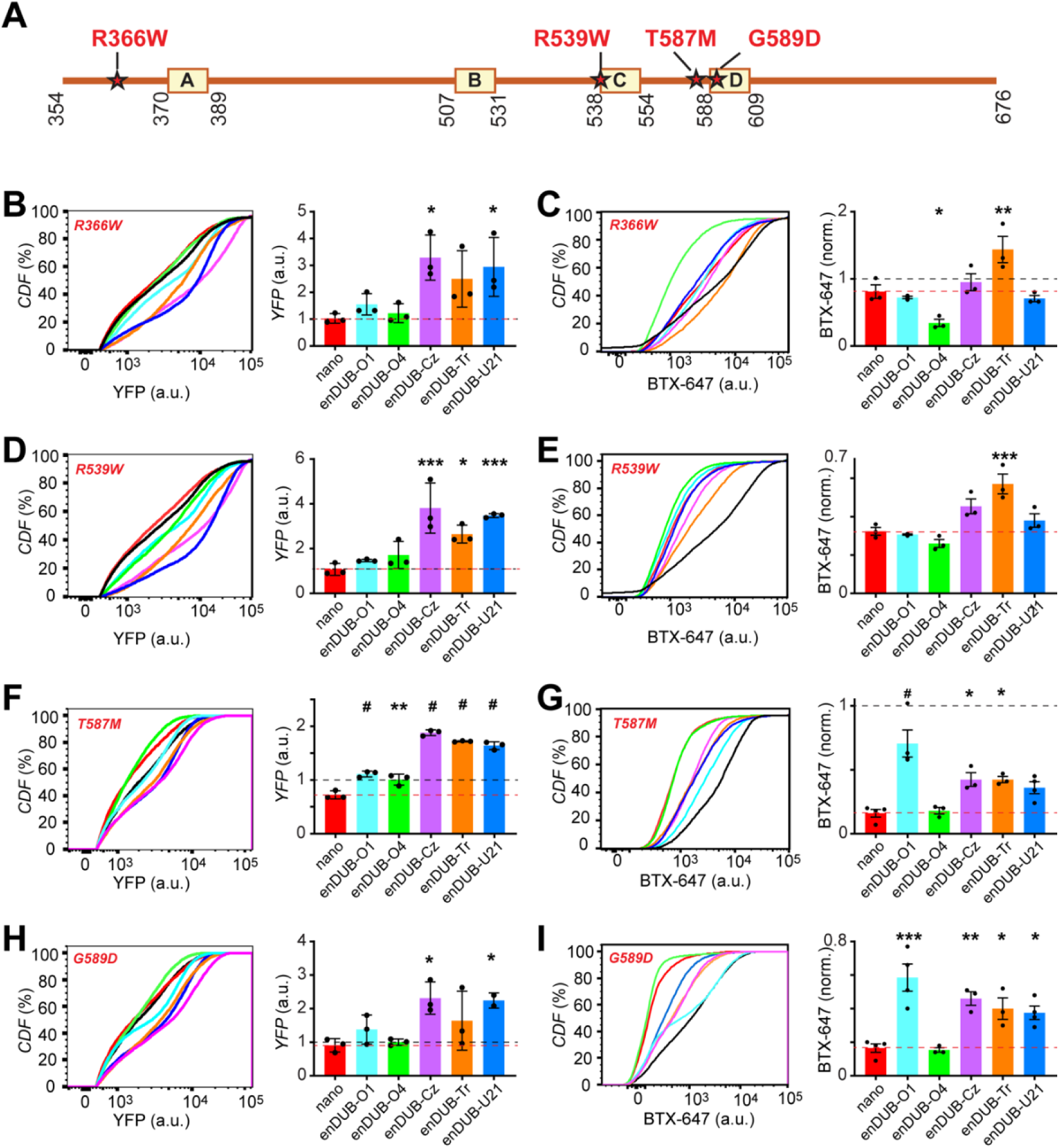
KCNQ1 disease mutants alter signature pattern of responsiveness to distinct enDUBs. (A) Scheme of LQT1 mutations mapped on a cartoon of the Q1 C-terminus. (B) Left, representative flow cytometry CDF plots of YFP fluorescence (channel total expression) in cells expressing BBS-KCNQ1[R366W]-YFP and either nano or an enDUBs. *Right*, Mean channel expression analyzed from YFP-and CFP-positive cells (*n*>5,000 cells per experiment, *N*=3). Data were normalized to the values from WT BBS-KCNQ1-YFP + nano control group (dotted line, black). Red dotted line corresponds to the values of mutant BBS-KCNQ1-YFP + nano. (C) *Left*, representative CDF plots showing BTX-647 fluorescence (surface channels) in cells expressing BBS-KCNQ1[R366W]-YFP and either nano or an enDUB. *Right*, mean channel expression analyzed from YFP-and CFP-positive cells (*n* > 5,000 cells per experiment, *N*=3). Data were normalized to values from WT BBS-Q1-YFP+nano control group (black dotted line). (D, E) Data for BBS-KCNQ1[R539W]-YFP; same format as B and C, respectively. (F, G) Data for BBS-KCNQ1[T587M]-YFP; same format as B and C, respectively. (H, I) Data for BBS-KCNQ1[G589D]-YFP; same format as B and C, respectively. *#p*<0.0001, ***p<0.001, **p<0.01 and *p<0.05; one-way ANOVA and Dunnett’s multiple comparisons test relative to the mutant values.

**Figure 9:**
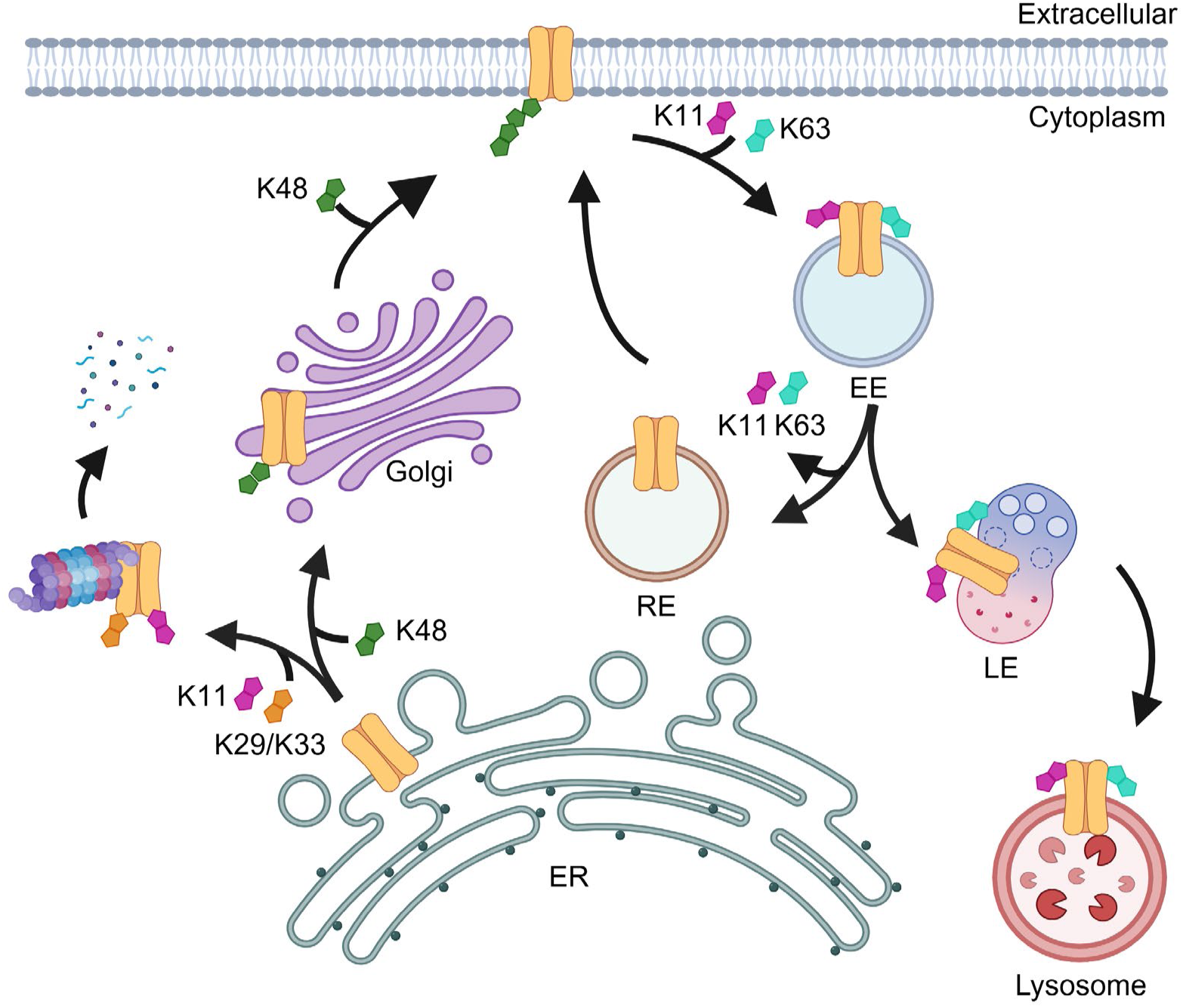
Differential roles of distinct polyubiquitin linkages in regulating KCNQ1 expression and trafficking in HEK293 cells. Cartoon showing proposed differential roles of distinct polyubiquitin chains in regulating KCNQ1 expression and trafficking among subcellular compartments in HEK 293 cells under basal conditions. This polyubiquitin code for KCNQ1 is not immutable and can be changed by both extrinsic (changing cellular conditions or cell types) and intrinsic (mutations in the substrate) factors.

Measuring surface density indicated the four mutations caused varying degrees of channel retention, ranging from moderate (R66W; 20% reduction) to intermediate (R539W; 60% reduction), and severe (T587M and G589D; 80% each) (Fig, 8C,E,G,I). The enDUBs had distinctive signature patterns on rescue of surface density: R366W surface density was increased by enDUB-Tr and decreased by enDUB-O4 (Fig. 8C); R539W was moderately rescued by enDUB-Tr and enDUB-Cz; T587M and G589D were most strongly rescued by enDUB-O1 with smaller effects produced by enDUB-Cz, enDUB-Tr, and enDUB-U21. Thus, these results reveal that missense mutations in KCNQ1 result in unique changes in polyubiquitin linkage code regulation of channel trafficking that potentially contributes to the molecular basis of disease.

## Discussion

Ubiquitination is an intricate posttranslational modification that regulates the functional expression of all proteins to control biology. The ubiquitin code refers to the complex system of ubiquitin writers, erasers, editors, and readers that govern the nature of ubiquitin signals on individual proteins and how they are decoded in cells to yield diverse functional outcomes (Dikic and Schulman, 2023; Komander and Rape, 2012; Kwon and Ciechanover, 2017). Understanding the ubiquitin code regulation of proteins is an active research frontier with broad implications for insights into physiology, disease mechanisms, and drug development. The prevalence of polyubiquitin chains with distinct linkages and topologies on proteins is a prominent aspect of the ubiquitin code (Akutsu et al., 2016; Komander and Rape, 2012; Kwon and Ciechanover, 2017; Yau and Rape, 2016). However, the functions of diverse polyubiquitin signatures on most proteins is unclear due to a lack of approaches to specifically tune polyubiquitin linkages on individual proteins in live cells. In this work we describe a method to address this core deficiency by developing linkage-selective enDUBs and apply them to investigate polyubiquitin regulation of KCNQ1-YFP expression, trafficking, and function in live cells. The results validate the utility of linkage-selective enDUBs to help decipher the polyubiquitin code regulation of protein function and expression in different cellular contexts, and provide new insights into divergent regulation of functional expression of an ion channel by distinct polyubiquitin linkages.

### Validity of enDUBs as a tool to decode polyubiquitin linkage function in cells

The strength of the conclusions drawn from the results obtained in this work hinges critically on the capacity of putative linkage-selective enDUBs to specifically hydrolyze intended polyubiquitin chains in a substrate selective manner. The approach was inspired by previous elegant work demonstrating the capacity of certain ovarian tumor (OTU) DUB catalytic domains to hydrolyze polyubiquitin chains in a linkage-selective manner *in vitro* (Mevissen et al., 2013; Ritorto et al., 2014). Nevertheless, a potential concern was that the linkage-selectivity of polyubiquitin chain hydrolysis would not hold *in situ*, or be compromised by prolonged tethering of DUB to substrate enabled by a nanobody. All four putative linkage-selective enDUBs (enDUB-O1, enDUB-O4, enDUB-Cz and enDUB-Tr) as well as the non-selective enDUB-U21 effectively reduced the amount of ubiquitin using a pan-ubiquitin antibody on KCNQ1-YFP. While reassuring in terms of confirming the *in vivo* enzymatic viability of the distinct enDUBs, this biochemical result in itself could not address whether the enDUBs were behaving in a linkage-selective manner.

Rather, their distinctive impact on KCNQ1-YFP expression, subcellular localization, and functional currents provides the strongest evidence that their enzymatic uniqueness is maintained *in situ*. A previous study examined the impact of K63 polyubiquitin linkages on post-endocytic sorting of epidermal growth factor receptor (EGFR) by fusing K63 linkage-selective deubiquitination enzyme AMSH (associated molecule with the Src homology 3 domain of signal transducing adaptor molecule) to the receptor. Mass spectrometry experiments suggested linkage-preference of AMSH for K63 over K48 hydrolysis was maintained even in this most extreme configuration where the DUB is covalently attached to the substrate (Huang et al., 2013). Our findings with the enDUBs are consistent with this previous result in terms of indicating that linkage-selective DUBs preserve their ubiquitin linkage preferences when tethered to a substrate. Nevertheless, enDUBs provide a clear advantage in versatility and generalizability as they can be adapted to use for any endogenous protein substrate without the need for covalent attachment of the DUB catalytic domain.

How do linkage-selective enDUBs contrast with prevailing methods to examine functions of distinct polyubiquitin linkages *in vivo*? A prevailing approach is to express K-to-R ubiquitin mutants in cells or organisms and observe the functional output. For example, this approach has been used effectively to examine global consequences of eliminating particular ubiquitin linkages in yeast (Xu et al., 2009), and to validate hypothesized functional roles on individual proteins in cells (Hoege et al., 2002; Jin et al., 2008; Lauwers et al., 2009). However, this approach alters ubiquitination of the entire cellular proteome in contrast to the substrate-specific nature of the linkage-selective enDUB method. Overall, the linkage-selective enDUB strategy uniquely fills a gap in the functional analyses of polyubiquitin linkages *in situ*, and is complementary to prevailing methods for addressing this aspect of the ubiquitin code (Sun and Zhang, 2022).

### Insights into polyubiquitin linkage regulation of KCNQ1 total expression

In HEK293 cells, KCNQ1-YFP expression was upregulated by enDUB-Cz, enDUB-Tr, and enDUB-U21, but not enDUB-O1 or enDUB-O4. These results indicate a dominant contribution of K11 and K29/K33 but not K63 or K48 to KCNQ1-YFP degradation in this cellular context. This result was unexpected because K48 was the most abundant linkage type associated with KCNQ1 and is the canonical polyubiquitin signal associated with protein degradation (Chau et al., 1989; Clague and Urbe, 2010; Komander and Rape, 2012; Kwon and Ciechanover, 2017). Precedence for K48 polyubiquitination not resulting in protein degradation is provided by the yeast transcription factor, Met4, which was found to encode a ubiquitin interaction motif (UIM) that intramolecularly binds K48 polyubiquitin and protects the protein from proteasomal degradation (Flick et al., 2006). Our results give reason to wonder whether KCNQ1 may similarly encode a UIM that protects the channel from K48 polyubiquitin-mediated proteasomal degradation. Future use of linkage-selective enDUBs may uncover that this phenomenon of non-degradative roles for K48 chains is more prevalent than currently appreciated.

Another surprise was the apparently dominant role of minor polyubiquitin chains associated with KCNQ1-YFP, K11 and K29/K33, in mediating channel degradation. Previous studies have linked both these chain types to protein degradation pathways; K11 chains assembled by the anaphase-promoting complex (APC/C) mediate degradation of cell cycle regulators (Jin et al., 2008; Peters, 2006), and K29 chains have also been associated with protein degradation (Johnson et al., 1995). Further, both K11-and K29-ubiquitin chains have been associated with endoplasmic reticulum-associated degradation (ERAD) (Locke et al., 2014; Lopata et al., 2020). The use of an apparently minor polyubiquitin species to control protein elimination may act as an effective valve to control the rate of channel degradation. A caveat is that our measure of the fraction of atypical polyubiquitin linkage types associated with the channel may be an underestimate because the mass spectrometry analyses was constrained to visually identified monomers and dimers of the channel from Coomassie-stained gels. Given that extensive polyubiquitination can result in substantial increases in molecular weight, the contribution of potential highly polyubiquitinated KCNQ1-YFP species would be underrepresented in the mass spectrometry data.

In sharp contrast to HEK293 cells, KCNQ1-YFP expression in adult guinea pig ventricular cardiomyocytes was robustly upregulated by enDUB-O1 and enDUB-U21, moderately by enDUB-Cz, and unaffected by enDUB-O4 and enDUB-Tr. These variations are consistent with fundamental differences in the ubiquitin code regulation of KCNQ1-YFP expression between HEK293 cells and cardiomyocytes. While K63 polyubiquitin is conventionally ascribed non-degradative functions, our results suggest a prominent role of K63 polyubiquitin in mediating degradation of KCNQ1-YFP in cardiomyocytes. One possibility is that lysosomal degradation of the channel plays a larger role in cardiomyocytes than we observed in HEK293 cells. A previous report indicated K63 seeds formation of K48/K63 branched ubiquitin chains that enable proteasomal degradation of certain substrates (Ohtake et al., 2018). While the lack of effect of enDUB-O4 on KCNQ1-YFP expression in cardiomyocytes does not support a role for K48 polyubiquitin in this context, it is possible that K63 may seed other types of branched ubiquitin chains that underlie degradation of the channel in heart cells.

### Insights into polyubiquitin linkage regulation of KCNQ1 subcellular localization and trafficking

The plasma membrane is the site of KCNQ1 function, making the impact of enDUBs on channel surface density particularly important. In HEK293 cells, the relative effects of the distinct enDUBs on KCNQ1 surface density varied substantially under different conditions, contrasting with the more stable impact on channel expression. EnDUB-O1 and enDUB-U21 had no effect on channel surface density under baseline conditions but effectively rescued KCNQ1 surface density that had been severely depressed by NEDD4-2 or ITCH; enDUB-Cz increased surface density under all conditions; and enDUB-O4 itself depressed KCNQ1 surface density under baseline conditions. Combined with the distinctive impact of these enDUBs on KCNQ1 subcellular localization and rates of forward trafficking and endocytosis, the results are consistent with K63, K11, and K29/K33 polyubiquitin chains mediating net intracellular retention of the channel, but achieved in different ways. K11 promotes ER retention and degradation, enhances endocytosis, and reduces channel recycling; K29/K33 primarily promotes ER retention and degradation; K63 enhances endocytosis and reduces channel recycling. Previous studies have reported roles of K63 polyubiquitin in mediating sorting of membrane proteins to post-endocytic vesicular compartments (Lauwers et al., 2009). To our knowledge, these studies provide the first evidence of roles for K11 and putatively K29/K33 polyubiquitin in controlling the subcellular localization and trafficking of an ion channel.

The pattern of enDUB effects on KCNQ1-YFP surface density under baseline conditions in cardiomyocytes differed from HEK293 cells. In cardiomyocytes, KCNQ1-YFP surface density was strongly enhanced by enDUB-O1; moderately increased by enDUB-Cz and enDUB-U21; unaffected by enDUB-Tr; and decreased by enDUB-O4. This signature pattern is closest to HEK293 cells over-expressing NEDD4-2 and may indicate that the ubiquitin code in cardiomyocytes under basal conditions is oriented towards having more K63 polyubiquitin on KCNQ1 compared to HEK293 cells.

Intriguingly, presumed removal of K48 by enDUB-O4 decreased KCNQ1-YFP surface density in both HEK293 cells and cardiomyocytes under baseline conditions. In HEK293 cells, this was accompanied by an increased retention in the ER and Golgi compartments. The most parsimonious interpretation of this result is that K48 polyubiquitin plays an active positive role in forward trafficking of KCNQ1 in both HEK293 cells and cardiomyocytes.

### Implications for disease and therapeutics development

Mutations in ion channels that impair their density at the cell surface underlie devastating diseases including cystic fibrosis, lethal cardiac arrhythmias, epilepsy, and intellectual disability. Trafficking of wild type ion channels can also be compromised in disease conditions such as diabetes in ways that contribute to pathology (Curran and Mohler, 2015; Harraz and Delpire, 2024). We previously showed that targeted deubiquitination with enDUB-O1 could rescue trafficking deficient mutations in KCNQ1 and cystic fibrosis transmembrane regulator (CFTR) that cause LQT1 and CF, respectively (Kanner et al., 2020). Consistent with the idea that net intracellular retention of ion channels due to mutations can occur in various ways we found that distinct KCNQ1 LQT1 mutations were differentially rescued by linkage-selective enDUBs. Surface density of R366W and R539W were most robustly increased by enDUB-Tr while T587M and G589D were most responsive to enDUB-O1. These results agree with previous findings that mutations in ion channels can alter their ubiquitination status in a manner that interferes with their functional expression (Aisenberg et al., 2022; Meacham et al., 2001; Okiyoneda et al., 2010; Wang et al., 2020). Our findings suggest that linkage-selective enDUBs could have utility in elucidating the precise trafficking deficits caused by distinct ion channel disease mutations, and in identifying the most efficacious polyubiquitin linkage type/s to target for rescue. Beyond enDUBs, divalent small molecule deubiquitinase targeting chimeras (DUBTACs) that recruit deubiquitinases to stabilize target proteins have also been developed (Henning et al., 2022). Deepened understanding of the versatility of the ubiquitin code combined with the uniqueness of distinct DUBs will expand the range of diseases and facilitate rational development of new therapeutics and tool compounds enabled by this induced proximity modality.

In summary, we have developed and applied linkage-selective enDUBs to dissect the role of distinct poly-ubiquitin chains in regulating KCNQ1 stability, subcellular trafficking, and function. The approach is modular and can be adapted to study the functions of diverse ubiquitin linkages on any intracellular protein with only the need to identify a suitable nanobody binder (or other antibody-mimetic) for the target. Nanobody binders can be readily generated by alpaca immunization and phage display technologies or by screening synthetic nanobody libraries (McMahon et al., 2018; Pardon et al., 2014). Alternatively, computational protein design algorithms may be used to design novel binders based on the sequence of the target protein (Watson et al., 2023). Future studies will expand the suite of linkage-selective enDUBs to include other linkage types. Altogether, linkage-selective enDUBs provide a unique tool that is complementary to prevailing methods to elucidate the intricate and dynamic polyubiquitin code *in vivo*.

## Supporting information

Compiled supplemental data

## Acknowledgements

We thank Ming Chen for technical support, and Drs. Arden Darko-Boateng and Yuki Utsugi for comments on the manuscript. This work was supported by grants RO1-HL121253, RO1-HL142111, R01-NS126850 and P01-HL164319 from the NIH (to HMC); AHA postdoctoral fellowship POST1019343 (to SSK); Medical Scientist Training Program grant (T32 GM007367) and NHLBI National Research Service Award (1F30-HL140878) (to SAK); EA was supported by NIGMS postdoctoral training grant T32 GM008464-32. Flow cytometry experiments were performed in the CCTI Flow Cytometry Core, supported in part by the NIH (S10RR027050). Confocal images were collected in the HICCC Confocal and Specialized Microscopy Shared Resource, supported by NIH (P30 CA013696).

## Author contributions

S.K.S. and S.A.K. designed and conducted experiments and analyzed data; X.Z., E.A., P.C., and R.S. conducted experiments and analyzed data; R.S.K. designed experiments and edited the paper; H.M.C. designed experiments and obtained funding; S.K.S and H.M.C. wrote the paper.

## Materials and Methods

### Plasmids and adenoviral generation

The construction of enDUB-O1 and enDUB-21 has been previously described (Kanner et al., 2020). To generate the other enDUBs we used polymerase chain reaction (PCR) to amplify the coding sequence for a GFP nanobody (vhhGFP4; nano) and cloned it into a xx-P2A-CFP cassette-containing mammalian expression vector (Kanner et al., 2017) using NheI and AflII sites. The catalytic domain of OTUD4 (residues 22-196) was amplified by PCR and cloned upstream of the GFP nanobody sequence to generate enDUB-O4-P2A-CFP. The sequences for the catalytic domains of Cezanne (residues 53-446) and TRABID (residues 245-696) were amplified by PCR and cloned downstream of GFP nanobody sequence to generate enDUB-Cz-P2A-CFP and enDUB-Tr-P2A-CFP, respectively. Catalytically inactive enDUBs were generated by site-directed mutagenesis using the QuikChange Lightning Site-Directed Mutagenesis kit (Stratagene) to introduce a point mutation at their catalytic cysteine residue.

KCNQ1 constructs were made as previously described (Aromolaran et al., 2014). Briefly, overlap extension PCR was used to fuse the sequence for enhanced yellow fluorescent protein (EYFP) in frame KCNQ1 C terminus. A 13-residue bungarotoxin binding site peptide (BBS) sequence was introduced between sequences for residues 148 and 149 in KCNQ1 extracellular S1–S2 loop using QuikChange Lightning Site-Directed Mutagenesis kit (Stratagene) according to the manufacturer’s instructions. LQT1 mutations were introduced in the N and C termini of KCNQ1 via site-directed mutagenesis as previously described (Aromolaran et al., 2014; Kanner et al., 2020). Plasmid vectors encoding NEDD4L (Addgene #27000) (Gao et al., 2009) and ITCH (Addgene #11427) (Magnifico et al., 2003) were gifts from Joan Massague and Allan Weismann, respectively.

Adenoviruses encoding BBS-KCNQ1-YFP, enDUB-O1-IRES-mCherry, enDUB-O4-IRES-mCherry, enDUB-Cz-IRES-mCherry, enDUB-Tr-IRES-mCherry, and enDUB-U21-IRES-mCherry were generated by Vector Builder.

### Cell culture and transfection

HEK293 and CHO cells were gifts from the laboratory of R. Kass (Columbia University). Cells were mycoplasma free, as determined by the MycoFluor Mycoplasma Detection kit (Invitrogen). Low-passage HEK293 cells were cultured at 37 °C in DMEM supplemented with 8% FBS and 100 mg/mL penicillin–streptomycin. For Western blot and flow cytometry experiments, transfection of HEK293 cells was accomplished using the calcium phosphate precipitation method. Briefly, plasmid DNA was mixed with 62 μl of 2.5 M CaCl2 and sterile deionized water (to a final volume of 500 μL). The mixture was added dropwise, with constant tapping, to 500 μL 2× HEPES-buffered saline containing (in mM) 50 HEPES, 280 NaCl and 1.5 Na2HPO4 (pH 7.09). The resulting DNA–calcium phosphate mixture was incubated for 20 min at room temperature (RT) and then added dropwise to HEK293 cells (60–80% confluent). Cells were washed with Ca^2+^-free PBS after 4–6 h and maintained in supplemented DMEM.

CHO cells were transiently transfected with desired constructs in 35-mm tissue culture dishes [KCNQ1 (0.5 μg) and nanobody–P2A– CFP (0.5 μg) or enDUB (0.5 μg)], in the presence or absence of E3 ligases (NEDD4L or ITCH; 0.5 μg) using X-tremeGENE HP (1:2 DNA:reagent ratio) according to the manufacturer’s instructions (Roche).

### Cardiomyocyte isolation and transduction

Isolation of adult guinea pig cardiomyocytes was performed in accordance with the guidelines of the Columbia University Animal Care and Use Committee. Before isolation, glass bottom culture dishes were precoated with 10 μg/mL laminin (Gibco). Adult Hartley guinea pigs (Charles River) were deeply anesthetized with 5% isoflurane and their hearts excised and mounted on a Langendorff perfusion apparatus. Ventricular myocytes were isolated by perfusing the heart first with KH solution (in mM, 118 NaCl, 4.8 KCl, 1 CaCl2, 25 HEPES, 1.25 K2HPO4, 1.25 MgSO4, 11 glucose, 0.02 EGTA, pH 7.4), followed by perfusion with enzymatic digestion solution containing 0.3 mg/mL Collagenase Type 4 (Worthington), 12.5 μg/mL protease, and 0.05% BSA in Ca^2+^-free KH buffer for 20 min. After digestion, 40 mL of a high-K^+^ solution containing120 mM potassium glutamate, 25 mM KCl, 10 mM HEPES, 1 mM MgCl2 and 0.02 mM EGTA, pH 7.4, was perfused through the heart. Cells were subsequently dispersed in the high-K^+^ solution. Healthy rod-shaped myocytes were cultured in Medium 199 (Life Technologies) supplemented with 10 mM HEPES (Gibco), 1×MEM non-essential amino acids (Gibco), 2 mM L-glutamine (Gibco), 20 mM D-glucose (Sigma-Aldrich), 1% (vol/vol) penicillin–streptomycin–glutamine (Fisher Scientific), 0.02 mg/mL vitamin B12 (Sigma-Aldrich) and 5% (vol/vol) FBS (Life Technologies) to promote attachment to dishes. After 3 h, the culture medium was changed to Medium 199 with 1% serum but otherwise supplemented as described above. Cultures were maintained in humidified incubators at 37 °C with 5% CO_2_. Adenoviral vectors were added to the medium and incubated overnight, followed by washing with fresh medium the next day. Experiments with cardiomyocytes were performed 1–2 d later.

### Flow Cytometry assays

Cell surface and total KCNQ1 channel expression were assayed by flow cytometry in live, transfected HEK293 cells as previously described (Aromolaran et al., 2014; Kanner et al., 2020; Zou et al., 2023). Briefly, 48 h post-transfection, cells cultured in 12-well plates were gently washed with ice cold PBS containing Ca^2+^ and Mg^2+^ (in mM: 0.9 CaCl_2_, 0.49 MgCl_2_, pH 7.4), and then incubated for 30 min in blocking medium (DMEM with 3% BSA) at 4°C. The cells were then incubated with 1 μM Alexa Fluor 647 conjugated α-bungarotoxin (BTX-647; Life Technologies) in DMEM/3% BSA on a rocker at 4°C for 1 h, followed by washing three times with PBS (containing Ca^2+^ and Mg^2+^). Cells were gently harvested in Ca^2+^-free PBS and assayed by flow cytometry using a BD LSRII Cell Analyzer (BD Biosciences, San Jose, CA, United States). CFP-and YFP-tagged proteins were excited at 407 and 488 nm, respectively, and Alexa Fluor 647 was excited at 633 nm. The optical pulse-chase assay was previously described in (Kanner et al., 2017; Zou et al., 2023). Briefly, post blocking, channels at the surface were first labeled with either α-bungarotoxin (Life Technologies) or biotin-conjugated α-bungarotoxin (Biotin-BTX; Life Technologies), before allowing for channel trafficking to resume for various times at 37 C, followed by labeling with 1 μM Alexa Fluor 647 conjugated α-bungarotoxin (BTX-647; Life Technologies) or 1 μM Alexa Fluor 647 conjugated Streptavidin (SA-647; Life Technologies) to measure channel forward and reverse trafficking, respectively.

### Electrophysiology

Patch clamp recordings of KCNQ1 or KCNQ1+KCNE1 currents (*I*_Ks_) were as previously described (Terrenoire et al., 2009; Zou et al., 2023). Cells plated in 3.5 cm culture dishes were mounted on the stage of an inverted microscope (OLYMPUS BH2-HLSH, Precision Micro Inc., Massapequa, NY). Whole-cell patch clamp currents were recorded at room temperature using an Axopatch 200B amplifier (Axon Instruments, Foster City, CA). Voltage-clamp pulse protocol consisted of a 500-ms step at a -70-mV holding potential a family of 2-s test pulse potentials (from -40 to +100 mV in 20-mV increments) and then a 2-s repolarizing pulses to -40 mV during which *I*_Ks_ tail current was measured. External solution contained: 132 mM NaCl, 4.8 mM KCl, 2 mM CaCl_2_, 1.2 mM MgCl_2_, 10 mM HEPES and 5 mM glucose (pH adjusted to 7.4 with NaOH). Internal solution contained: 110 mM KCl, 5 mM ATP-K_2_, 11 mM EGTA, 10 mM HEPES, 1 mM CaCl_2_ and 1 mM MgCl_2_ (pH adjusted to 7.3 with KOH). Pipette series resistance was typically 1.5 -3 MΩ when filled with internal solution. Currents were sampled at 10 kHz and filtered at 5 kHz. Traces were acquired at a repetition interval of 10 seconds.

### Confocal imaging

At 24 hr post-transfection, HEK cells were split onto fibronectin-coated 8-well glass bottom chamber slides (Ibidi GmbH). In some conditions, cells were treated overnight with 5 µM MG132 or 100 µM Chloroquine to block the proteosome or lysosome, respectively. The next day, cells were treated with 100 µM cycloheximide for 1h and fixed with 4% paraformaldehyde (PFA) for 10 mins at room temperature (RT). Cells were washed three times with PBS and permeabilized for 10 mins with 0.2% Triton X-100 in PBS with 0.2 M glycine and 3% BSA. The cells were washed thrice with PBS and incubated 30 mins in blocking buffer (PBS with 0.2 M glycine and 3% BSA), followed by overnight incubation with antibodies targeting subcellular organelles [Calnexin (Ab22595), RCAS1 (D2B6N), Rab9A (D52G8), EEA1 (C45B10) and LAMP2 (L0668)]. The cells were then washed thrice with PBS and incubated for 1h with HRP-conjugated goat anti-rabbit secondary antibody (1:5000) and imaged with a Nikon A1RMP confocal microscope using a 60x oil-immersion objective.

Cardiomyocytes were plated onto 35-mm MatTek dishes (MatTek Corporation) and fixed with 4% formaldehyde for 10 min at RT. Cells were stained with BTX-647 as described above. Images were captured on a Nikon A1RMP confocal microscope with a 40x oil-immersion objective.

### Immunoprecipitation, Western blot, and proteomics sample preparation

HEK293/CHO cells were washed once with Ca^2+^-free PBS, harvested, and resuspended in RIPA lysis buffer containing (in mM): Tris (20, pH 7.4), EDTA (1), NaCl (150), 0.1% (wt/vol) SDS, 1% Triton X-100, 1% sodium deoxycholate and supplemented with protease inhibitor mixture (10 μL/mL, Sigma-Aldrich, St. Louis, MO), PMSF (1 mM, Sigma-Aldrich), N-ethylmaleimide (2 mM, Sigma-Aldrich) and PR-619 deubiquitinase inhibitor (50 μM, LifeSensors). Lysates were prepared by incubation at 4°C for 1 hr, with occasional vortexing, and cleared by centrifugation (10,000 × g, 10 min, 4°C). Supernatants were transferred to new tubes, with aliquots removed for quantification of total protein concentration using the BCA protein estimation kit (Pierce Technologies).

For immunoprecipitation, lysates were pre-cleared by incubation with 20 µL Protein A/G Sepharose beads (Rockland) bound to anti-rabbit IgG (Sigma) for 3h at 4°C. Equivalent total protein amounts were added to spin-columns containing 50 µL Protein A/G Sepharose beads incubated with 3 µg anti-KCNQ1 (Alomone, Jerusalem, Israel), and tumbling overnight at 4°C. Immunoprecipitates were washed 3–5 times with RIPA buffer, spun down at 500 × g, eluted with 40 µL of warmed sample buffer [50 mM Tris, 10% (vol/vol) glycerol, 2% SDS, 100 mM DTT, and 0.2 mg/mL bromophenol blue], and boiled (55°C, 15 min). Proteins were resolved on a 4–12% Bis·Tris gradient precast gel (Life Technologies) in MOPS-SDS running buffer (Life Technologies) at 200 V constant for ∼1 hr. We loaded 10 μL of the PageRuler Plus Prestained Protein Ladder (10–250 kDa, Thermo Fisher, Waltham, MA) alongside the samples.

For proteomic analysis, the gels were stained with SimplyBlue (ThermoFisher Scientific) and KCNQ1 monomer and dimer bands were excised. In-gel digestion was performed as previously described (Shevchenko et al., 2006), with minor modifications. Gel slices were washed with 1:1 Acetonitrile and 100 mM ammonium bicarbonate for 30 min then dehydrated with 100% acetonitrile for 10 min until shrunk. The excess acetonitrile was removed, gel slices were dried in speed-vacuum at room temperature for 10 minutes and then reduced with 5 mM DTT for 30 min at 56°C in an air thermostat, cooled down to room temperature, and alkylated with 11 mM IAA for 30 min with no light. Gel slices were then washed with 100 mM of ammonium bicarbonate and 100 % acetonitrile for 10 min each. Excess acetonitrile was removed and dried in a speed-vacuum for 10 min at room temperature and the gel slices were re-hydrated in a solution of 25 ng/μL trypsin in 50 mM ammonium bicarbonate for 30 min on ice and digested overnight at 37°C in an air thermostat. Digested peptides were collected and further extracted from gel slices in extraction buffer (1:2 ratio by volume of 5% formic acid: acetonitrile) at high speed, shaking in an air thermostat. The supernatants from both extractions were combined and dried in a speed-vacuum. Peptides were dissolved in 3% acetonitrile/0.1% formic acid.

For Western blotting, protein bands were transferred from the gel by tank transfer onto a nitrocellulose or Immobilon-P PVDF membrane (Sigma-Aldrich) (3.5 hr, 4°C, 30 V constant) in transfer buffer (25 mM Tris pH 8.3, 192 mM glycine, 15% (vol/vol) methanol, and 0.1% SDS). The membranes were blocked with a solution of 5% nonfat milk (BioRad) in tris-buffered saline-tween (TBS-T) (25 mM Tris pH 7.4, 150 mM NaCl, and 0.1% Tween-20) for 1 hr at RT and then incubated overnight at 4°C with one of the primary antibodies (anti-KCNQ1, Alomone, APC-022, 1:1,000 or anti-actin antibody (1:2000, Abcam, USA) in blocking solution. The blots were washed with TBS-T three times for 10 min each and then incubated with secondary horseradish peroxidase-conjugated antibody for 1 hr at RT. After washing in TBS-T, the blots were developed with a chemiluminescent detection kit (Pierce Technologies) and then visualized on a gel imager. Membranes were then stripped with harsh stripping buffer (2% SDS, 62 mM Tris pH 6.8, 0.8% ß-mercaptoethanol) at 50°C for 30 min, rinsed under running water for 2 min, and washed with TBST (3x, 10 min). Membranes were blocked with 5% milk again and re-blotted with anti-ubiquitin (LifeSensors, VU1, 1:500).

### Liquid chromatography with tandem mass spectrometry (LC-MS/MS)

Desalted peptides were injected in an EASY-SprayTM PepMapTM RSLC C18 50cm X 75cm ID column (Thermo Scientific) connected to an Orbitrap FusionTM TribridTM (Thermo Scientific). Peptides elution and separation were achieved at a non-linear flow rate of 250 nL/min using a gradient of 5%-30% of buffer B (0.1% (v/v) formic acid, 100% acetonitrile) for 110 minutes with a temperature of the column maintained at 50°C during the entire experiment. The Thermo Scientific Orbitrap Fusion Tribrid mass spectrometer was used for peptide tandem mass spectroscopy (MS/MS). Survey scans of peptide precursors are performed from 350 to 1500 m/z at 120K full width at half maximum (FWHM) resolution (at 200 m/z) with a 2 x 105 ion count target and a maximum injection time of 60 ms. The instrument was set to run in top speed mode with 3-second cycles for the survey and the MS/MS scans. After a survey scan, MS/MS was performed on the most abundant precursors, i.e., those exhibiting a charge state from 2 to 6 of greater than 5 x 10^3^ intensity, by isolating them in the quadrupole at 1.6 Th. We used Higher-energy C-trap dissociation (HCD) with 30% collision energy and detected the resulting fragments with the rapid scan rate in the ion trap. The automatic gain control (AGC) target for MS/MS was set to 5 x 10^4^ and the maximum injection time was limited to 30 ms. The dynamic exclusion was set to 30 s with a 10 ppm mass tolerance around the precursor and its isotopes. Monoisotopic precursor selection was enabled.

### Data and statistical analysis

Patch clamp data, shown as mean ± S.E., were acquired using pCLAMP 8.0 (Axon Instruments) and analyzed with Origin 7.0 (OriginLab, Northampton, MA) and Clampfit 8.2 (Axon Instruments). Flow cytometry data were analyzed using FlowJo 10.8 software. Raw mass spectrometric data were analyzed using the Proteome Discoverer 2.4 to perform database search and LFQ quantification at default settings. PD2.4 was set up to search with the reference human proteome database downloaded from UniProt and performed the search trypsin digestion with up to 2 missed cleavages. Peptide and protein false discovery rates (FDR) were all set to 1%. The following modifications were used for protein identification and LFQ quantification: Carbamidomethyl(C) was set as fixed modification and variable modifications of Oxidation (M) and Acetyl (Protein N-term), DiGly (K) and deamination for asparagine or glutamine (NQ). Results obtained from PD2.4 were further used to quantify relative diGly peptide abundances under the different conditions and identifying sites modified on KCNQ1. Statistical data analysis was assessed with Student’s *t*-test for comparison between two groups and one-way ANOVA for comparisons among more than two groups. In the Figures, statistically significant differences from control values are marked by asterisks (*, p<0.05; **, p<0.01; ***, p<0.001; #, p<0.0001).

## Material availability statement

Plasmid constructs for non-commercial purposes can be obtained by request from the corresponding author after publication of the manuscript.

---

**Supplemental Figure 1: KCNQ1-YFP subcellular distribution under proteosomal and lysosomal inhibition.**

(A) Representative confocal images of HEK293 cells that were treated with MG132 expressing KCNQ1-YFP (green) and immunolabelled subcellular organelles (red -ER, Golgi, early endosomes (EE), late endosomes (LE) and lysosomes). (B) Representative confocal images of HEK293 cells that were treated with chloroquine expressing KCNQ1-YFP (green) and immunolabelled subcellular organelles (red - ER, Golgi, EE, LE and lysosomes). (C) Pearson’s correlational coefficient of KCNQ1-YFP with the subcellular organelles ER (*n*>7; \**p*<0.05, one-way ANOVA with Dunnett’s multiple comparisons test), Golgi (*n*>6; *ns* one-way ANOVA with Dunnett’s multiple comparisons test), EE (*n*>7; *ns*, one-way ANOVA), LE (*n*>8, *ns*, one-way ANOVA) and lysosomes (*n*>8;\*\*\**p*<0.001, one-way ANOVA with Dunnett’s multiple comparisons test).

**Supplemental Figure 2: Impact of MG132 and chloroquine on total and surface expression of KCNQ1-YFP**

(A) Representative flow cytometry CDF plot showing surface (BTX_647_) fluorescence (left) and total (YFP) fluorescence in cells expressing KCNQ1-YFP under untreated (black) and chloroquine treated (gray) conditions. (B) Quantification of flow cytometry experiments for KCNQ1-YFP surface expression (BTX-647 fluorescence) analyzed from YFP-positive cells in untreated (black), MG132 (red) and chloroquine (gray) treated conditions (*n*>5,000 cells per experiments; *N*=5-6; \**p*<0.05, one-way ANOVA). Data were normalized to the control group. (C) Quantification of flow cytometry experiments for KCNQ1-YFP total expression (YFP fluorescence) analyzed from YFP-positive cells in untreated (black), MG132 (red) and chloroquine (gray) treated conditions (*n*>5,000 cells per experiments; *N*=5-6; *ns, p*>0.05, one-way ANOVA). Data were normalized to the control group.

**Supplemental Figure 3: Exemplar ms2 spectra**

(A) Exemplar ms2 spectra traces for K11 and (B) K27 ubiquitin digly peptides.

**Supplemental Figure 4: Design of linkage-selective and non-specific enDUBs** Schematic of enDUB design.

**Supplemental Figure 5: Catalytic activity of the enDUBs is essential for its impact on KCNQ1-YFP.** Representative blot of immunoprecipitated KCNQ1-YFP co-expressed with the enDUBs and probed with anti-KCNQ1 (left). The same blot was stripped and probed again with anti-ubiquitin (right)

**Supplemental Figure 6: Targeting the DUBs with GFP nanobody is essential for its impact on KCNQ1-YFP.**

(A) Scheme. KCNQ1 co-expressed with the enDUBs. (B) Quantification of flow cytometry experiments for KCNQ1 surface expression analyzed from CFP-positive cells (n > 5,000 cells per experiments; *N*=2; *ns*, one-way ANOVA). Data were normalized to the values from the nano control group (dotted line).

**Supplemental Figure 7: Confocal images of subcellular redistribution of KCNQ1-YFP by the enDUBs.**

Exemplar confocal images of HEK293 cells expressing KCNQ1-YFP (green) and enDUBs with immunostaining of the subcellular organelles ER, golgi, EE, LE and lysosome.

**Supplemental Figure 8: Differential impact of the enDUBs on KCNQ1-YFP surface density in the presence of Rab11DN.**

(A) Cartoon of impact of Rab11DN mediated recycling of KCNQ1-YFP. (B) Quantification of flow cytometry experiments for KCNQ1 surface expression analyzed from YFP-positive cells (n>5,000 cells per experiments; *N*=3; \*\*\**p*<0.001, \*\**p*<0.001 and \**p*<0.05, one-way ANOVA with Dunnett’s multiple comparisons). Data were normalized to the values from the nano control group (dotted line).

**Supplemental Figure 9: Differential effect of NEDD4L and ITCH on PY mutant KCNQ1.**

(A) Representative flow cytometry CDF plot showing surface (BTX_647_) fluorescence in cells expressing Y662-KCNQ1-YFP NEDD4L and ITCH. Quantification of flow cytometry experiments for KCNQ1-YFP surface expression analyzed from YFP-positive cells (n>5,000 cells per experiments; *N*=3). Data were normalized to the values from the nano control group (dotted line). (B) Exemplar Y662A KCNQ1-YFP current traces from whole-cell patch clamp measurements in CHO cells with NEDD4L (orange) and ITCH (cyan).

**Supplemental Figure 10: KCNQ1 sequence.**

Sequence of KCNQ1 channel. Cytosolic regions are highlighted in green, Transmembrane segments are in blue, extracellular regions are in gray, PY motif is in maroon. Cytosolically accessible unmodified and modified (as detected by mass spectrometry) lysine (K) residues are highlighted in pink and in yellow, respectively.

