## Supplementary material for "Decoding polyubiquitin regulation of K_V_7. 1 functional expression with engineered linkage-selective deubiquitinases": Compiled supplemental data

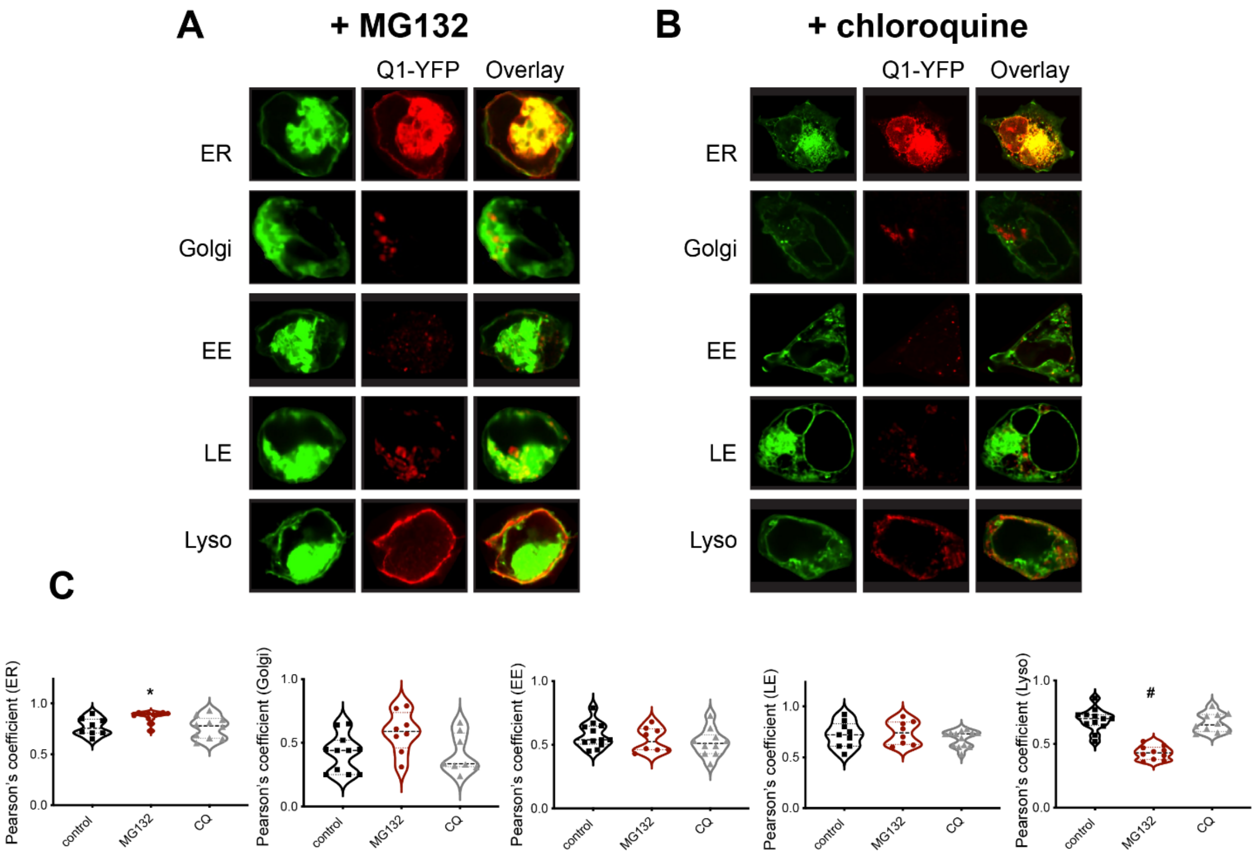


**Supplemental Figure 1: KCNQ1-YFP subcellular distribution under proteosomal and lysosomal inhibition.**

(A) Representative confocal images of HEK293 cells that were treated with MG132 expressing KCNQ1-YFP (green) and immunolabelled subcellular organelles (red - ER, Golgi, early endosomes (EE), late endosomes (LE) and lysosomes). (B) Representative confocal images of HEK293 cells that were treated with chloroquine expressing KCNQ1-YFP (green) and immunolabelled subcellular organelles (red - ER, Golgi, EE, LE and lysosomes). (C) Pearson's correlational coefficient of KCNQ1-YFP with the subcellular organelles ER ($n>7$; *$p<0.05$, one-way ANOVA with Dunnett's multiple comparisons test), Golgi ($n>6$; *ns* one-way ANOVA with Dunnett's multiple comparisons test), EE ($n>7$; *ns*, one-way ANOVA), LE ($n>8$, *ns*, one-way ANOVA) and lysosomes ($n>8$; ***$p<0.001$, one-way ANOVA with Dunnett's multiple comparisons test).

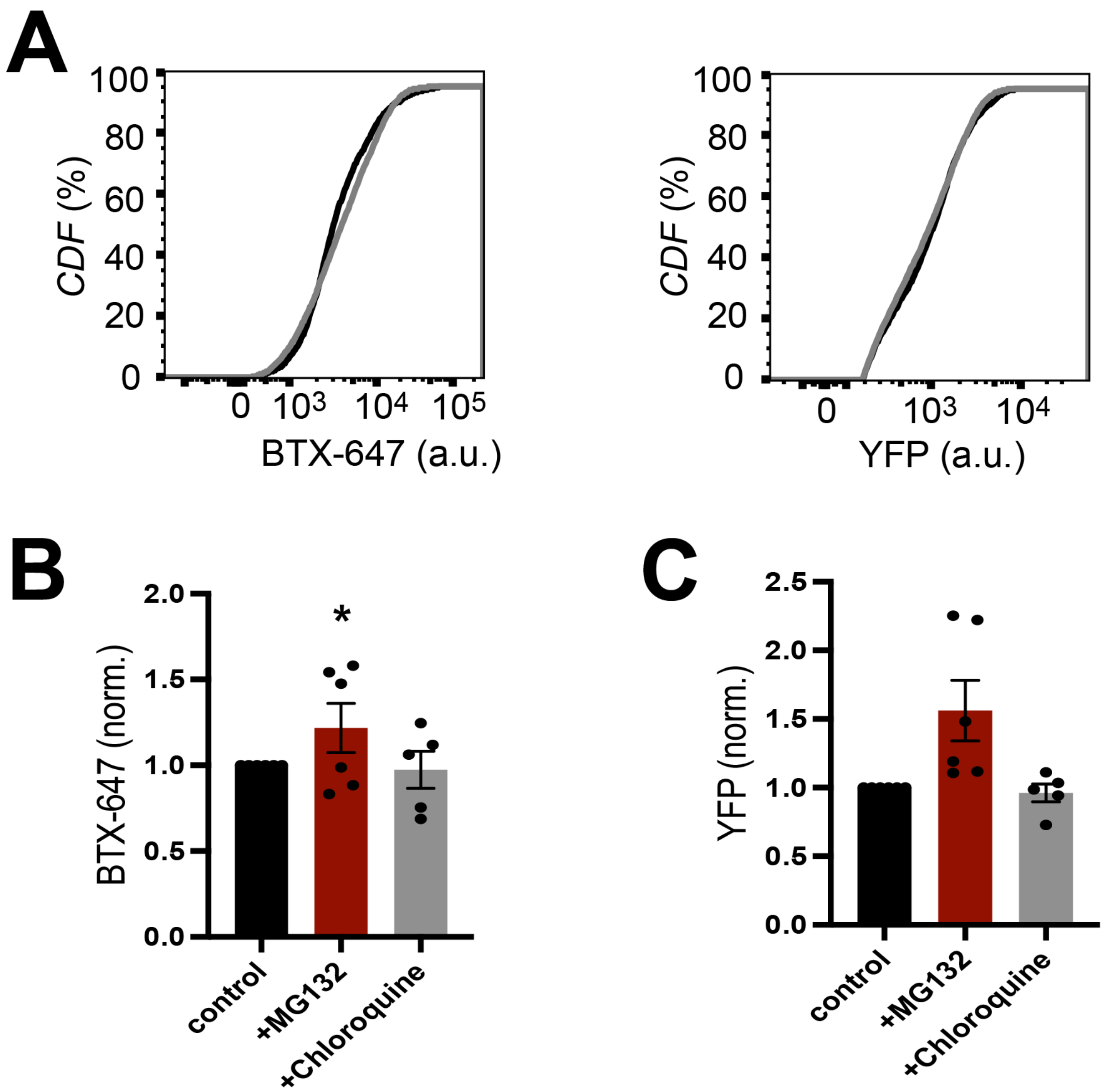


**Supplemental Figure 2: Impact of MG132 and chloroquine on total and surface expression of KCNQ1-YFP**

(A) Representative flow cytometry CDF plot showing surface ($BTX_{647}$) fluorescence (left) and total (YFP) fluorescence in cells expressing KCNQ1-YFP under untreated (black) and chloroquine treated (gray) conditions. (B) Quantification of flow cytometry experiments for KCNQ1-YFP surface expression (BTX-647 fluorescence) analyzed from YFP- positive cells in untreated (black), MG132 (red) and chloroquine (gray) treated conditions ($n$>5,000 cells per experiments; $N$=5-6; *$p$<0.05, one-way ANOVA). Data were normalized to the control group. (C) Quantification of flow cytometry experiments for KCNQ1-YFP total expression (YFP fluorescence) analyzed from YFP- positive cells in untreated (black), MG132 (red) and chloroquine (gray) treated conditions ($n$>5,000 cells per experiments; $N$=5-6; *ns*, $p$>0.05, one-way ANOVA). Data were normalized to the control group.

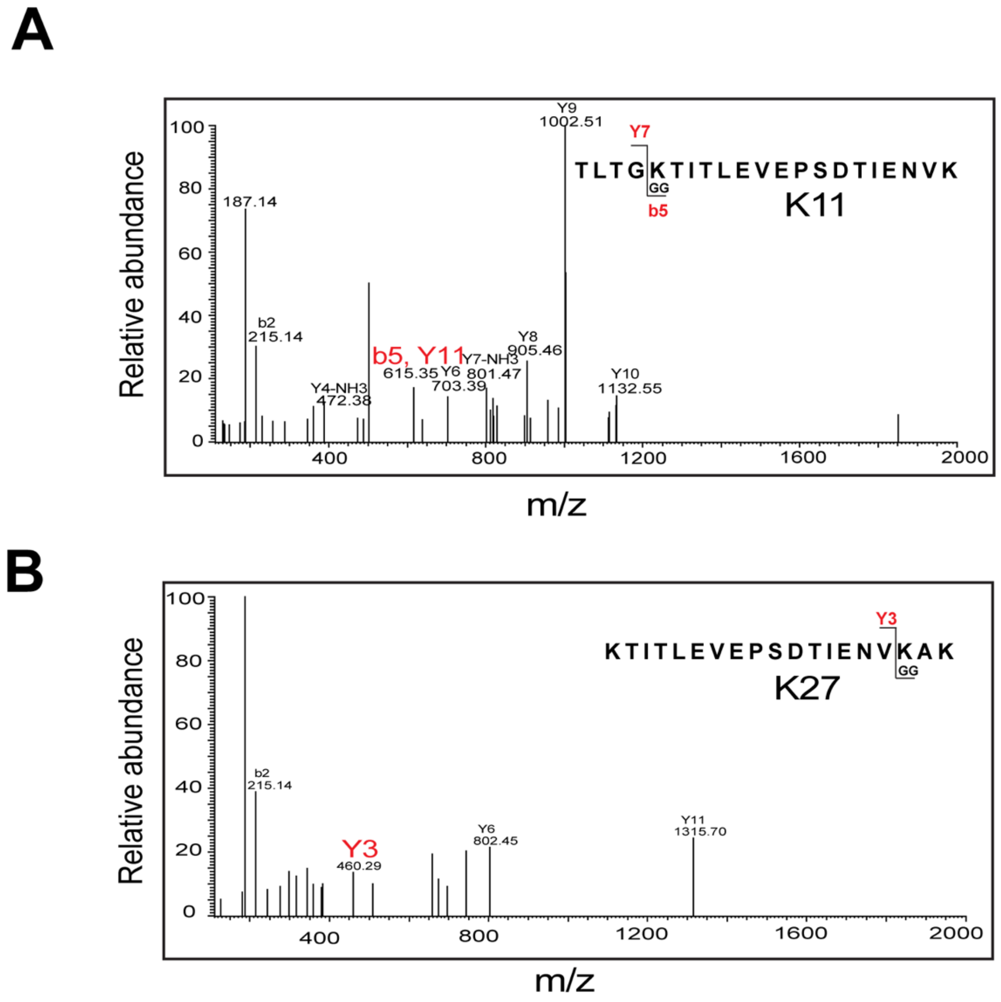


**Supplemental Figure 3: Exemplar ms2 spectra**

(A) Exemplar ms2 spectra traces for K11 and (B) K27 ubiquitin digly peptides.

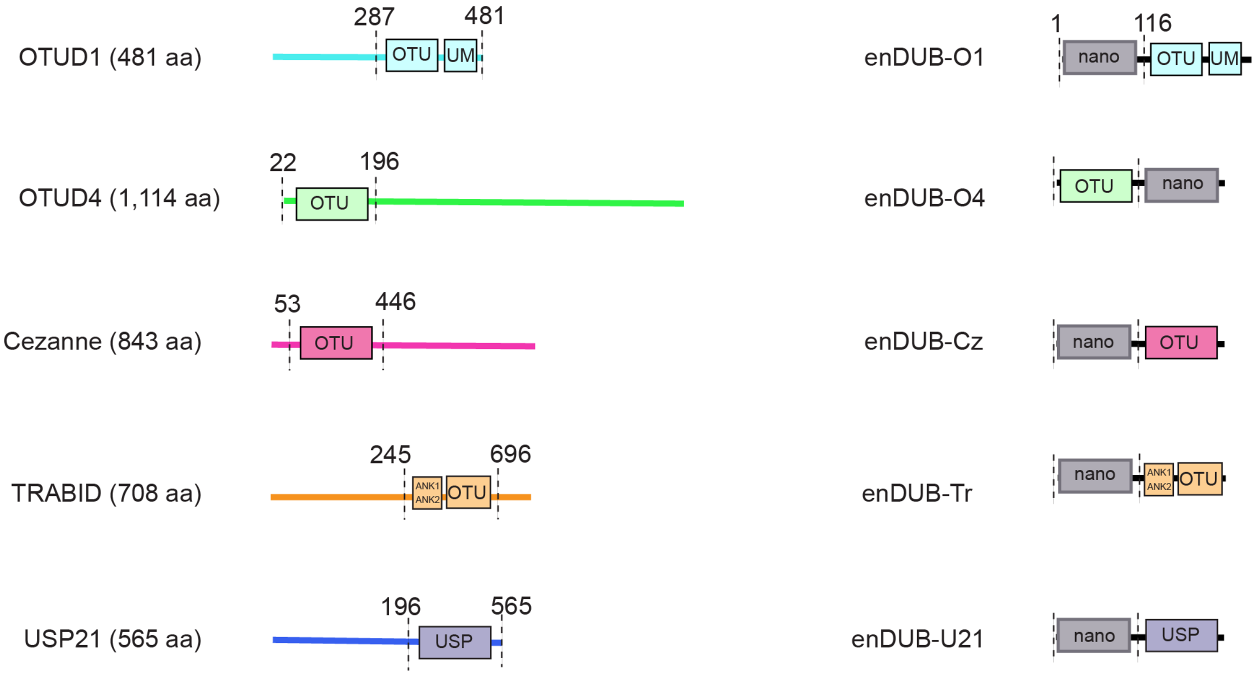


**Supplemental Figure 4: Design of linkage-selective and non-specific enDUBs**

Schematic of enDUB design.

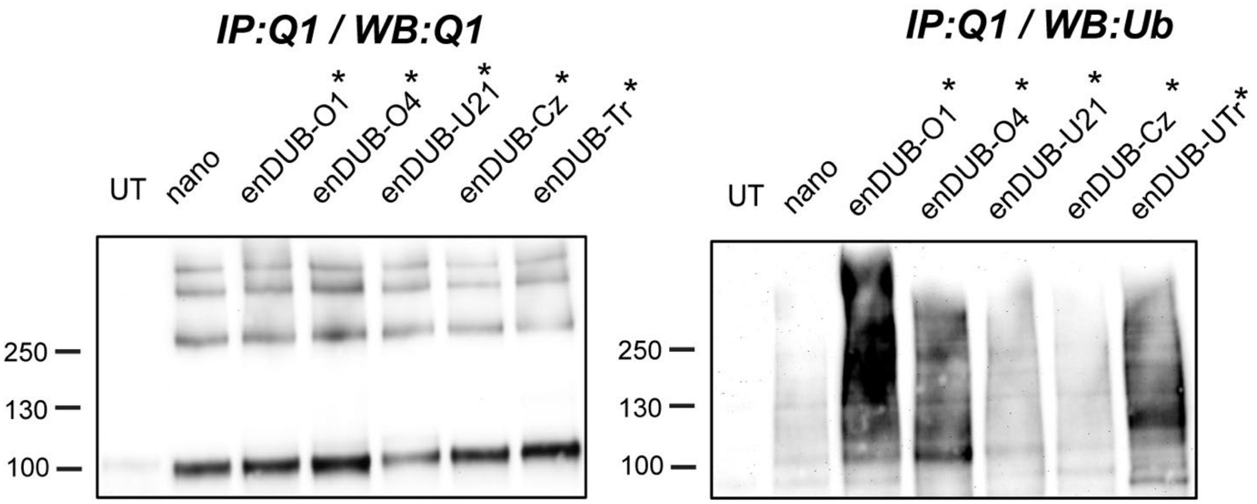


**Supplemental Figure 5: Catalytic activity of the enDUBs is essential for its impact on KCNQ1-YFP.**

Representative blot of immunoprecipitated KCNQ1-YFP co-expressed with the enDUBs and probed with anti-KCNQ1 (left). The same blot was stripped and probed again with anti-ubiquitin (right)

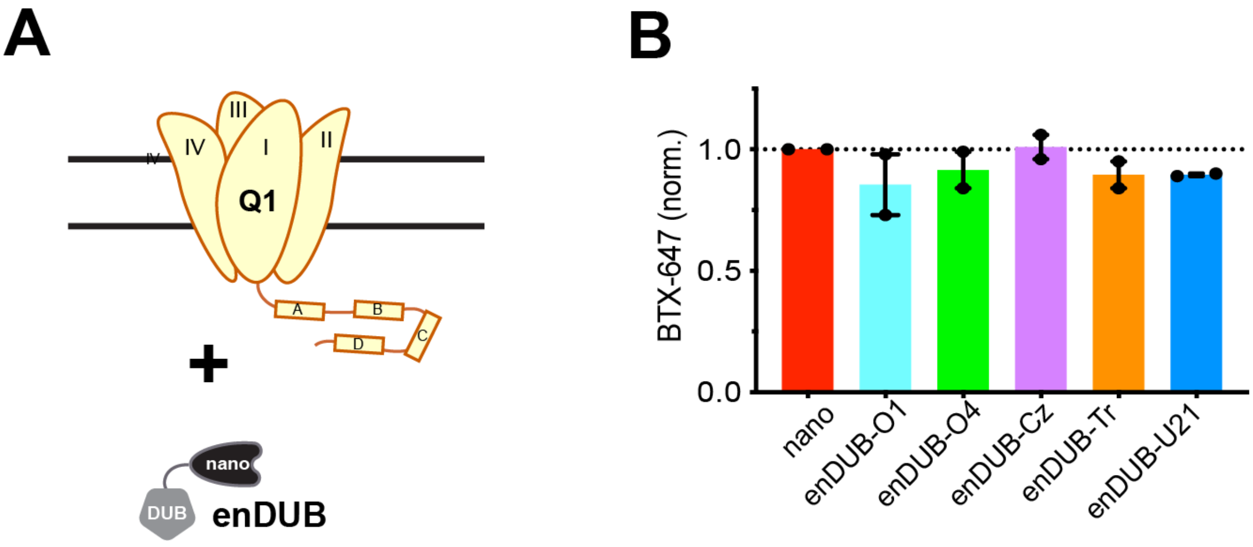


**Supplemental Figure 6: Targeting the DUBs with GFP nanobody is essential for its impact on KCNQ1-YFP.**

(A) Scheme. KCNQ1 co-expressed with the enDUBs. (B) Quantification of flow cytometry experiments for KCNQ1 surface expression analyzed from CFP- positive cells (n > 5,000 cells per experiments; *N*=2; *ns*, one-way ANOVA). Data were normalized to the values from the nano control group (dotted line).

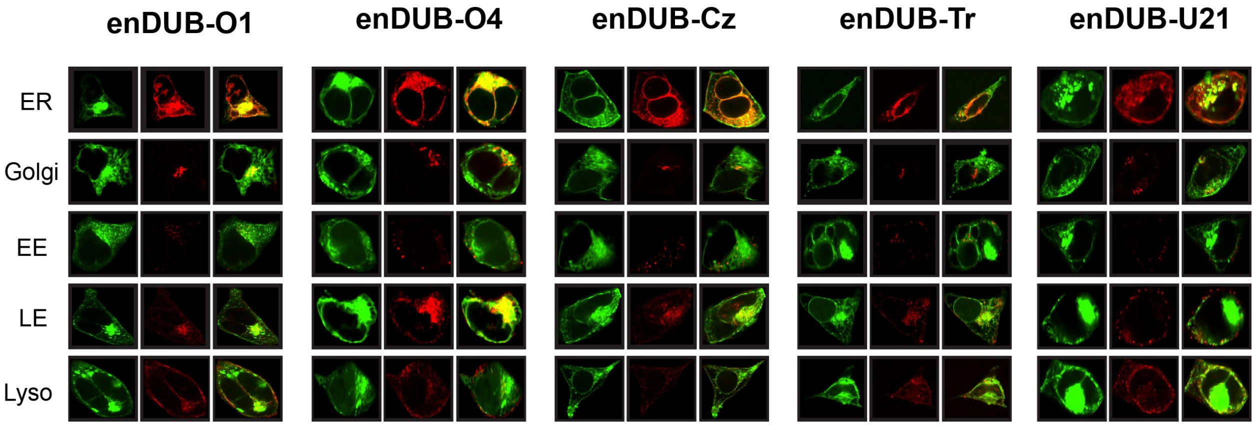


**Supplemental Figure 7: Confocal images of subcellular redistribution of KCNQ1-YFP by the enDUBs.**

Exemplar confocal images of HEK293 cells expressing KCNQ1-YFP (green) and enDUBs with immunostaining of the subcellular organelles ER, golgi, EE, LE and lysosome.

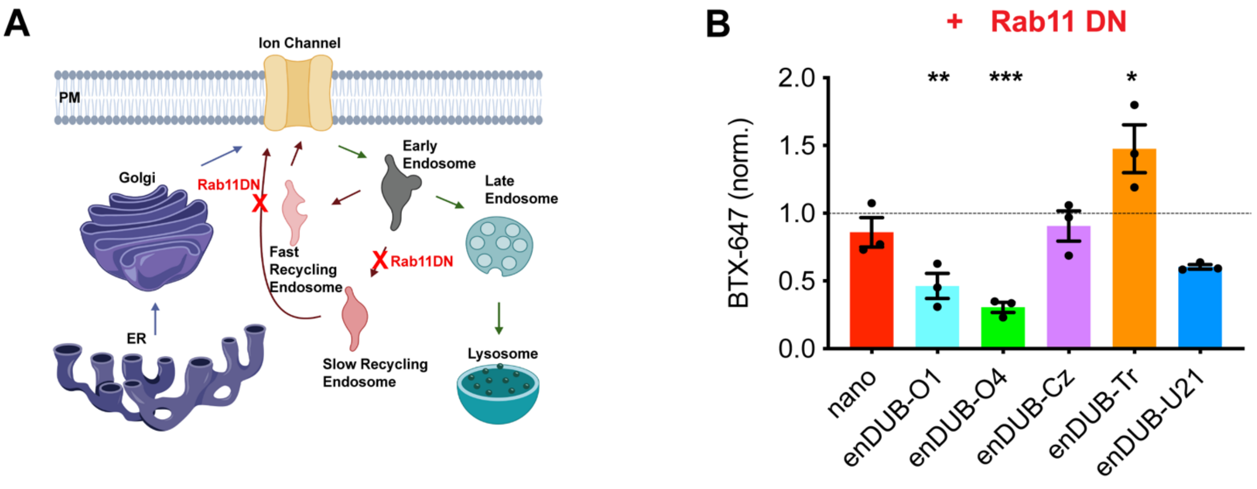


**Supplemental Figure 8: Differential impact of the enDUBs on KCNQ1-YFP surface density in the presence of Rab11DN.**

(A) Cartoon of impact of Rab11DN mediated recycling of KCNQ1-YFP. (B) Quantification of flow cytometry experiments for KCNQ1 surface expression analyzed from YFP- positive cells (n>5,000 cells per experiments; *N*=3; ***$p$<0.001, **$p$<0.001 and *$p$<0.05, one-way ANOVA with Dunnett's multiple comparisons). Data were normalized to the values from the nano control group (dotted line).

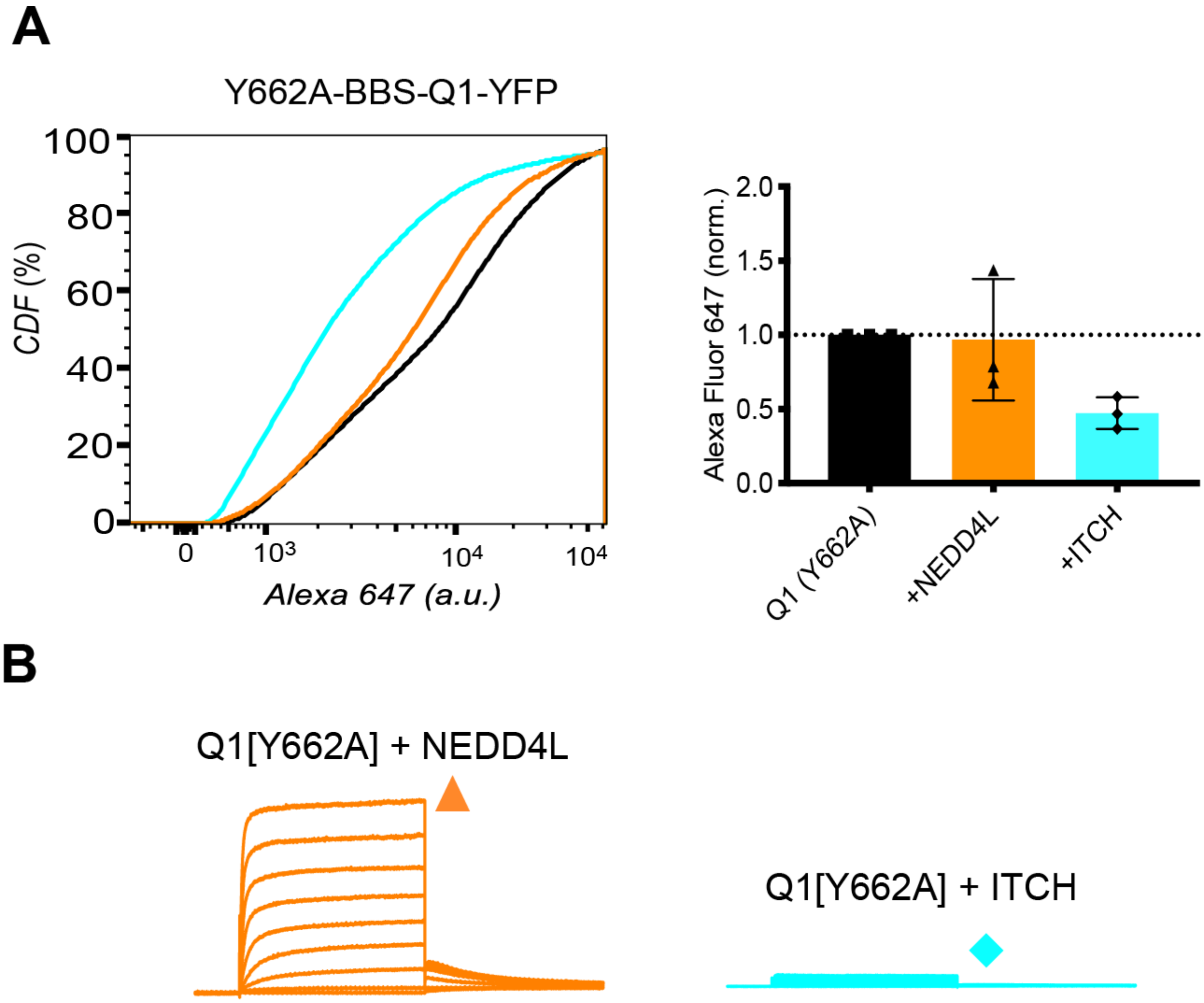


**Supplemental Figure 9: Differential effect of NEDD4L and ITCH on PY mutant KCNQ1.**

(A) Representative flow cytometry CDF plot showing surface ($BTX_{647}$) fluorescence in cells expressing Y662-KCNQ1-YFP NEDD4L and ITCH. Quantification of flow cytometry experiments for KCNQ1-YFP surface expression analyzed from YFP- positive cells (n>5,000 cells per experiments; *N*=3). Data were normalized to the values from the nano control group (dotted line). (B) Exemplar Y662A KCNQ1-YFP current traces from whole-cell patch clamp measurements in CHO cells with NEDD4L (orange) and ITCH (cyan).

MAAASSPPRAERKRWGWGRLPGARRGSAGLAKKCPFSLELAEG
GPAGGALYAPIAPGAPGPAPPASPAAPAAPPVASDLGPRPPVS
LDPRVSIYSTRRPVLARTHVQGRVYNFLERPTGWKCFVYHFAV
FLIVLVCLIFSVLSTIEQYAALATGTLFWMEIVLVVFFGTEYV
VRLWSAGCRSKYVGLWGRLRFARKPISIIDLIVVVASMVVLCV
GSKGQVFATSAIRGIRFLQILRMLHVDRQGGTWRLLGSVVFIH
RQELITTLYIGFLGLIFSSYFVYLAEKDAVNESGRVEFGSYAD
ALWWGVVTVTTIGYGDKVPQTWVGKTIASCFSVFAISFFALPA
GILGSGFALKVQQKQRQKHFNRQIPAAASLIQTAWRCYAAENP
DSSTWKIYIRKAPRSHTLLSPSPKPKKSVVVKKKKFKLDKDNG
VTPGEKMLTVPHITCDPPEERRLDHFSVDGYDSSVRKSPTLLE
VSMPHFMRTNSFAEDLDLEGETLLTPITHISQLREHHRATIKV
IRRMQYFVAKKKFQQARKPYDVRDVIEQYSQGHLNLMVRIKEL
QRRLDQSIGKPSLFISVSEKSKDRGSNTIGARLNRVEDKVTQL
DQRLALITDMLHQLLSLHGGSTPGSGGPPREGGAHITQPCGSG
GSVDPELFLPSNTLPTYEQLTVPRRGPDEGS

**Supplemental Figure 10: KCNQ1 sequence.**

Sequence of KCNQ1 channel. Cytosolic regions are highlighted in green, Transmembranal segments are in blue, extracellular regions are in gray, PY motif is in maroon. Cytosolically accessible unmodified and modified (as detected by mass spectrometry) lysine (K) residues are highlighted in pink and in yellow, respectively.
